# Genetic and transcriptomic characteristics of RhlR-dependent quorum sensing in cystic fibrosis isolates of *Pseudomonas aeruginosa*

**DOI:** 10.1101/2021.12.21.470774

**Authors:** Kyle L. Asfahl, Nicole E. Smalley, Alexandria Chang, Ajai A. Dandekar

## Abstract

In people with the genetic disease cystic fibrosis (CF), bacterial infections involving the opportunistic pathogen *Pseudomonas aeruginosa* are a significant cause of morbidity and mortality. *P. aeruginosa* uses a cell-cell signaling mechanism called quorum sensing (QS) to regulate many virulence functions. One type of QS consists of acyl-homoserine lactone (AHL) signals produced by LuxI-type signal synthases, which bind a cognate LuxR-type transcription factor. In laboratory strains and conditions, *P. aeruginosa* employs two AHL synthase/receptor pairs arranged in a hierarchy, with the LasI/R system controlling the RhlI/R system and many downstream virulence factors. However, *P. aeruginosa* isolates with inactivating mutations in *lasR* are frequently isolated from chronic CF infections. We and others have shown that these isolates frequently use RhlR as the primary QS regulator. RhlR is rarely mutated in CF and environmental settings. We were interested if there were reproducible genetic characteristics of these isolates and if there was a central group of genes regulated by RhlR in all isolates. We examined five isolates and found signatures of adaptation common to CF isolates. We did not identify a common genetic mechanism to explain the switch from Las-to Rhl-dominated QS. We describe a core RhlR regulon encompassing 20 genes encoding 7 products. These results suggest a key group of QS-regulated factors important for pathogenesis of chronic infection, and position RhlR as a target for anti-QS therapeutics. Our work underscores the need to sample a diversity of isolates to understanding QS beyond what has been described in laboratory strains.

## Introduction

Persistent infection with the environmental bacterium and opportunistic pathogen *Pseudomonas aeruginosa* is a common clinical problem. Burn wounds, indwelling catheters, and the airways of people with the genetic disease cystic fibrosis (CF) all provide an ecological niche suited to *P. aeruginosa* infection [1, 2]. Acute infections begin with motile *P. aeruginosa* expressing a variety of secreted toxins, proteases, and virulence factors to establish infection, aided by intrinsic resistance to several classes of antibiotics [3]. A transition follows, where *P. aeruginosa* adapts to favor a biofilm lifestyle that protects the pathogen from host immune factors [3–5]. Increased host inflammation and fluid viscosity further aid biofilm construction, particularly in the CF lung, where *P. aeruginosa* infection is associated with morbidity and mortality [6, 7]. Ultimately, *P. aeruginosa* strains adapted to chronic infections persist despite aggressive treatment.

A substantial fraction of the virulence factor program of *P. aeruginosa* is regulated via a cell-cell communication system termed quorum sensing (QS), where a diffusible signal binds a cognate receptor that is a transcriptional activator, thereby allowing monitoring of population density and a coordinated transcriptional response [8, 9]. QS provides a mechanism for *P. aeruginosa* to specifically modulate virulence gene expression based on environmental constraints within the host [8, 10]. *P. aeruginosa* has two complete QS systems that use acyl-homoserine lactone (AHL) signals: LasI synthesizes *N*-3-oxo-dodecanoyl-homoserine lactone (3OC12-HSL), which is bound by LasR; and RhlI synthesizes *N*-butanoyl homoserine lactone (C4-HSL), which is bound by RhlR (Figure 1) [11]. A third QS circuit in *P. aeruginosa* uses alkyl-quinolone (AQ) signals that are bound by the receptor and LysR-type transcription factor PqsR (also called MvfR) and termed the *Pseudomonas* quinolone signal (PQS) system [12]. In laboratory strains, the Rhl and PQS systems are regulated by LasR, and together these *P. aeruginosa* QS systems allow concerted population-wide regulation of more than 200 genes in response to cell density [13, 14].

**Figure 1.**
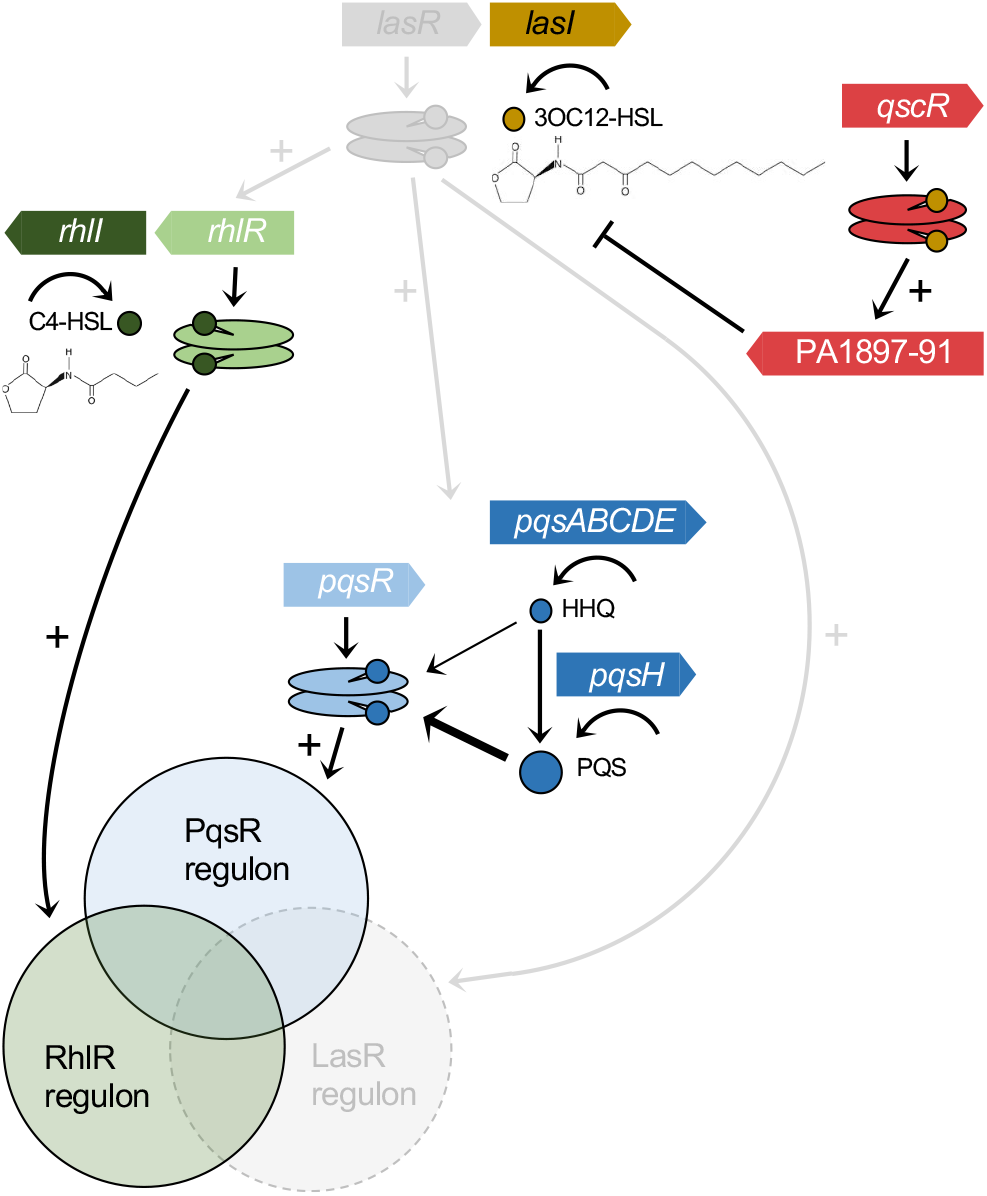
Las-independent RhlR-QS circuitry in *P. aeruginosa*. In laboratory strains, LasR activation is required for downstream activation of the RhlR-QS (green) and PQS (blue) circuitry. The CF isolates analyzed in this study harbor deleterious mutations that inactivate LasR (gray components), yet still use RhlR and the PQS system to regulate overlapping sets of target genes. The orphan regulator QscR (red) binds the LasI-generated signal and activates a single linked operon.

*A* complex interplay of selective pressures contributes to diversification of *P. aeruginosa* during the shift from acute to chronic infection, and the QS systems of *P. aeruginosa* appear to be common targets of selection. The gene encoding LasR is often mutated in isolates from chronic CF infections, yet mutations in RhlR are rare [15–18]. This observation suggested that RhlR may function independently of LasR in some isolates, in contrast to the LasR-dominated circuitry exhibited in laboratory strains (Figure 1). This notion is supported through observations of isolates with variant LasR alleles from the lungs of CF patients, where Rhl-QS appears to regulate production of elastase, rhamnolipid surfactants, and the toxic secondary metabolite pyocyanin [17].

We are interested in understanding the evolution of *P. aeruginosa* QS as strains become adapted to the CF lung and switch from LasR-to RhlR-dominated QS. The identification of such isolates also presents an opportunity: the hierarchical arrangement of QS in laboratory strains, where LasR is required for activation of Rhl-QS under most circumstances, has presented a roadblock to understanding the RhlR regulon. The scope of Rhl-QS-mediated gene regulation has been approached previously via transcriptome analysis of laboratory strains with mutations in *rhlR* or *rhlI*, through supplementation of synthetic C4-HSL signals, or both [19, 20]. These studies identified lists of genes that are generally considered to be subsets of QS targets regulated by LasR, and because LasR activation was instrumental in these designs, understanding the explicit importance of RhlR in their regulation remained elusive. We hypothesized that a core regulon for RhlR exists among isolates with a common mode of infection, that of the CF lung, and that a common genomic adaptation might explain the switch to Las-independent Rhl-dominated QS.

Here, we used a collection of CF isolates with natural inactivating mutations in LasR to determine a core regulon for the QS receptor RhlR among 5 infection isolates. We show that these isolates do not respond to the 3OC12-HSL signal through LasR, but are still able to use the Rhl-QS system to activate RhlR target promoters in a density-dependent manner, control pyocyanin production, and augment extracellular proteolysis. We conducted a comparative genomic analysis to find an identifiable switch that could yield Las-independent RhlR-QS. We used deletion of *rhlR* and supplementation of synthetic QS signals to yield 5 parallel RhlR-off and RhlR-on conditions for use in an RNA-seq-based RhlR transcriptome analysis for each isolate. We found that among 227 genes regulated by RhlR in at least 1 isolate in our cohort, just 20 genes comprise the core RhlR regulon.

## Results

### Isolate selection

We identified CF isolates with naturally occurring *lasR* variant alleles through a screen of a collection of >2500 *P. aeruginosa* strains isolated from the lungs of children enrolled in the Early *Pseudomonas* Infection Control (EPIC) observational study[21]. Our group has previously reported on the prevalence of nonsynonymous *lasR* variants in this collection (~22%) with an initial characterization of the QS activity in 31 isolates harboring unique *lasR* variant alleles[17].

For the present study we sought to understand Las-independent RhlR-QS, so our initial screen criteria required the isolates to have loss-of-function *lasR* alleles while also having positive phenotype for C4-HSL production in our previous study[17]. Of the isolates we were interested in, we required that the isolates accept transcriptional fusion reporter plasmids and be amenable to genetic manipulation. We selected 5 such isolates, all of which were from different patients (Table 1).

**Table 1.** CF isolate characteristics.

| Isolate | C4-HSL (uM)<sup>1</sup> | LasR mutation |  |
| --- | --- | --- | --- |
|  |  | Nucleotide | Amino acid |
| E104 | 17.73 | A532G | T178A |
| E113 | 3.45 | T55C | W19R |
| E125 | 13.43 | C280T | Q94Stop |
| E131 | 26.54 | A580G | S194G |
| E167 | 7.44 | G339del | Frameshift |

<sup>1</sup>C4-HSL determinations from Feltner et al. 2016.

### RhlR controls gene promoters and phenotypes associated with virulence in a cohort of CF isolates

First, we queried whether QS transcriptional activity was in fact independent of LasR using transcriptional fusion reporter plasmids. We asked if the *lasR* alleles in our CF isolates retained LasR activity. To do so, we measured GFP fluorescence from cells transformed with a P*_lasI_-gfp* reporter. The *lasI* promoter is LasR-activated and, consistent with the idea that the mutations in *lasR* were inactivating, we found fluorescence was significantly lower in each of our clinical isolates than in PAO1. Because lack of gene activation could reflect inadequate signal concentrations rather than nonfunctional protein, we also asked if addition of the LasR signal 3OC12-HSL increased fluorescence in any of the isolates (Figure 2A). It did not.

**Figure 2.**
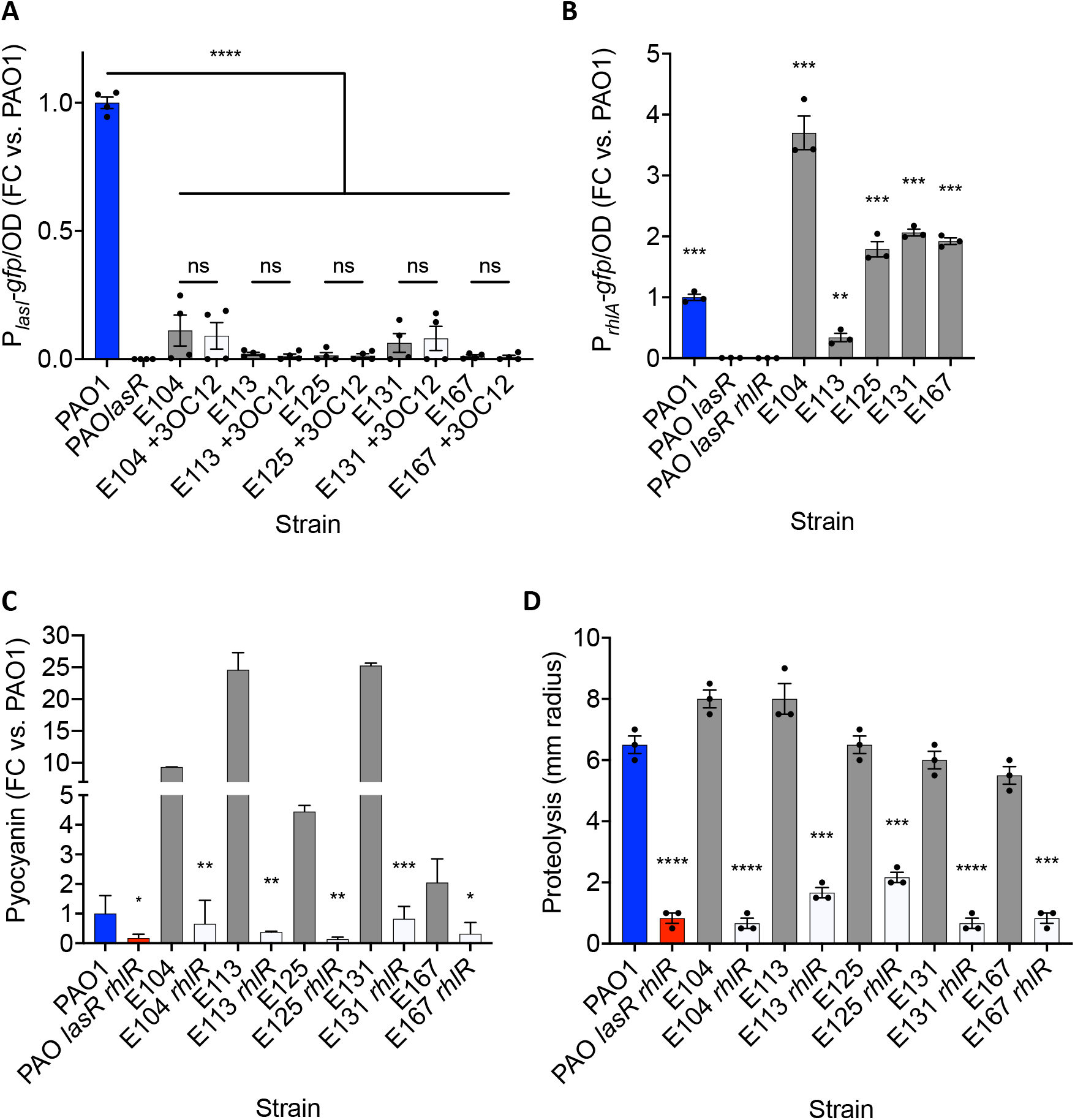
Las-independent QS transcription and phenotypes. (A) pP*_lasI_-gfp* reporter activity, normalized to OD_600_ and scaled to PAO1. 3OC12-HSL was added to a final concentration of 2 μM. (B) pP*_rhlA_-gfp* reporter activity, normalized to OD_600_ and scaled to PAO1. (C) Pyocyanin production after 18 h, normalized to OD_600_ and scaled to PAO1. (D) Extracellular proteolysis, determined as the zone of clearing on skim-milk agar after 48 h. For all panels means, standard error, and the results of unpaired *t* tests are shown (*n*≥3). *, *p* < 0.05; **, *p* < 0.01; ***, *p* < 0.001; ****, *p* < 0.0001.

Next, to measure transcription from a RhlR-regulated promoter, we measured GFP fluorescence using a P*_rhlA_-gfp* reporter. The promoter for *rhlA* is strictly controlled by RhlR[22, 23]. Despite mutations in *lasR* in our CF isolates, each strain showed activation of the *rhlA* promoter (P*_rhlA_-gfp*) that was significantly greater than that of the *lasR* mutant laboratory strain PAO1 (Figure 2B). We then queried whether the timing of induction of the *rhlA* promoter was advanced compared to PAO1 by following activation over time; no strain was significantly advanced in P*_rhlA_-gfp* induction (Figure S1). Together these results indicated that in these CF isolates LasR is inactive and unresponsive to its signal, but RhlR remains active as a QS transcriptional regulator.

In an effort to understand the range of QS-controlled phenotypes in our CF isolate cohort, we examined production of the redox-active pigment and secreted toxin pyocyanin, and extracellular proteolysis. First, we determined the level of pyocyanin produced by our isolates and their isogenic *rhlR* mutants. Production of pyocyanin was elevated in every isolate compared to the laboratory strain PAO1, ranging from roughly 2-fold that of PAO1 in strain E167 up to 25-fold in strain E131 (Figure 2C). Regardless of the level produced by each wildtype isolate, deletion of *rhlR* in each isolate background reduced pyocyanin to levels lower than that of wildtype PAO1, and similar to that of the reference QS double deletion mutant PAO1 *lasR rhlR*. We then expanded our analysis to extracellular proteolysis, using skim-milk agar plates[24]. In the laboratory strain PAO1, secreted proteases such as the QS-controlled factor LasB elastase diffuse away from the colony, producing a zone of reduced opacity with a radius that varies with regulation of protease. QS mutants of PAO1 show a marked reduction in this radius of clearing (Figure 2D). We found that while the radius of clearing varied between individual isolates and relative to PAO1, deletion of *rhlR* significantly reduced this radius in each isolate (Figure 2D). Together, these phenotypes indicated that despite harboring inactivating mutations in *lasR*, these isolates still use RhlR to regulate production of virulence factors.

### Genomic analysis

We sequenced and assembled complete, closed genomes of each isolate for comparative genomic analysis and for use as strain-specific transcriptomic mapping references. Genome sizes were all greater than that of PAO1 (6.26 Mbp), ranging from 6.67 Mbp for strain E131 to 6.89 Mbp for strain E125 (Table S1). The number of annotated features was also greater in each isolate than PAO1, with an average of 658 additional annotated features (Table S1). Strain E167 also harbored a stably maintained plasmid of approximately 47 kbp.

In an effort to understand the relatedness of our CF isolate cohort to other sequenced isolates of *P. aeruginosa*, we compared our isolate genomes to a curated collection of genomes on the publicly available IMG/MER database[25]. We constructed a phylogeny of our isolates (5), strain E90 from our previous study[26], all publicly available complete genomes associated with cystic fibrosis (10) or airway isolation (5), and the laboratory strains PAO1 and PA14 (21 strains total; Figure 3). We found our 5 isolates were distributed throughout the phylogeny. Isolates E113 and E131 were closely clustered with the epidemic CF infection strains described from Liverpool (LESB58) and Manchester (C3719), with our previously described strain E90, and laboratory strains PAO1 and PA14. On the other hand, isolates E104, E125, and E167 were more distantly related and separated from the phylogeny, similar to the Danish transmissible strain DK2 and the composite assembly strain PACS2, isolated from a 6-month-old with CF. Together, our phylogenetic analysis indicated our 5-isolate cohort was genomically diverse and represented a variety of CF isolate origins.

**Figure 3.**
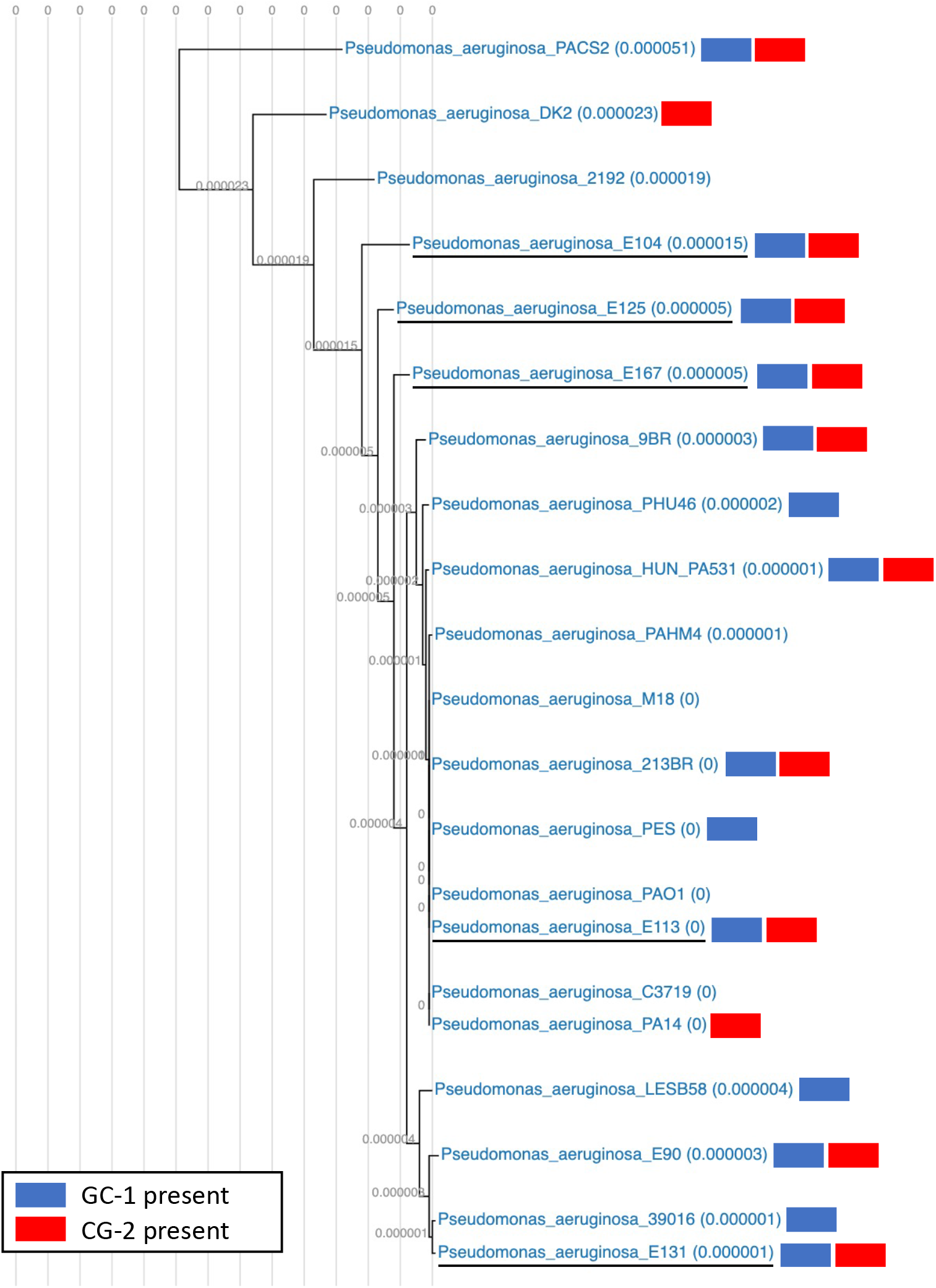
Phylogenetic distribution of CF isolate cohort. CF isolates in this study (underlined) are well distributed in a phylogeny that includes laboratory strains and previously published genomes of *P. aeruginosa* isolates with airway or CF infection origins. Blue and red denote the presence of gene cassette 1 and 2, respectively (GC-1, GC-2), in the genomes of individual isolates.

Previous investigations have indicated the genome of *P. aeruginosa* may experience purifying selection during adaptation to the CF lung, with selection against certain acute virulence factors [15, 27]. To gain an understanding of how the content of our cohort isolate genomes compare to both PAO1 and previously published genomic analyses of CF isolates, we conducted a pangenomic analysis. We found a total of 7156 genes in the pangenome of our isolate cohort and PAO1, with a core genome of 5353 genes present in all six strains. We found deletions or inactivating mutations in 197 genes common amongst the isolate cohort but normally present in laboratory strains (Table S2). 74 of these genes were missing entirely from all 5 genomes. This group of commonly mutated or missing genes includes 16 genes annotated as transcriptional regulators, including *lasR*. No other gene with an established direct regulatory link to the Las, Rhl, or PQS QS systems was found to be mutated. Mutation of other transcriptional regulators such as *mexT* may provide a pathway to active RhlR QS in the absence of a functional LasR, a feature observed during *in vitro* evolution of PAO1 [28, 29], but we found no evidence of mutations in *mexT* in our study.

Other regulators found to be missing or mutated included those involved in amino acid catabolism (*amaR*, *dguR*), terpene utilization (*atuR*), and multidrug efflux and antibiotic susceptibility (*armR*, *oprR*, *cmrA*). The list of commonly lost or mutated genes also included other genes known to be targeted by selection during long-term evolution in the CF lung[15, 18, 30]: O-antigen biosynthesis (*hisF2H2*, *wzx*, *wzy*), T3SS (*exsD*), exotoxin A (*xqhB*), and twitching motility (*pilA*). Genes involved in interspecific interactions with neighboring microbes were also mutated in all 5 isolates: T2SS (*xqhB*), T4SS (*pilB/S/Y1*, *fimT*), T6SS (*tse6*, *tssC1/L1*), and the S-pyocin killing mechanism (*pyoS5, pys2*, PA3866).

In addition to genes commonly lost or mutated relative to PAO1, we found 112 genes in common among our cohort that are not present in the laboratory strain (Table S3). 87% (98) of these common non-PAO1 genes are organized into cassettes that harbor annotations for integrative transfer (e.g. *pil* genes), along with some strain-specific gene content. These consist of two organized and conserved cassettes of 28 and 70 genes (gene cassette-1 and −2, GC-1 and GC-2). Both cassettes are widespread within the strains of our analyzed phylogeny (Figure 3). GC-1 includes genes coding for at least 3 putative regulators, mitomycin antibiotic biosynthesis proteins, a collection of acyl-CoA modifying enzymes and several hypothetical proteins. GC-2 harbors at least 15 IncI1-group conjugative plasmid-associated genes (PFGI-1-like, *parB, pilL/N/P/Q/S/U/M* are present), genes coding for 2 putative regulators, the mercuric resistance protein MerT, genes involved in T4SS (*virB4*), a T3SS effector Hop protein, and several hypothetical proteins. Beyond these conserved cassettes, the list of commonly acquired genes includes additional genes involved in T4SS (*virD4*), heavy metal resistance (copper efflux protein *cusA* and a lead/cadmium/zinc/ mercury ATPase), an RpoS-like sigma factor, and numerous genes related to mobile genetic elements. Finally, the stably-maintained plasmid in strain E167 (47.4 kbp) harbors genes encoding a HigB/A toxin-antitoxin system, in addition to numerous hypothetical proteins and plasmid maintenance genes.

### A core regulon for RhlR

A primary goal of the present study was to discern commonalities among RhlR QS gene regulation in CF-adapted *P. aeruginosa*. To this end, we evaluated the transcriptomes of each isolate at the entry to stationary phase (OD_600_ 2.0) (Figure 4A), using differential expression (DE) analysis between wild-type CF isolates and their isogenic *rhlR* deletion mutants. This a point in growth previous studies have shown to provide a reasonable census of QS activation among various strains [19, 31]. We also added signals to the wild-type condition to avoid differences in gene regulation due to variable signal production (Table 1).

**Figure 4.**
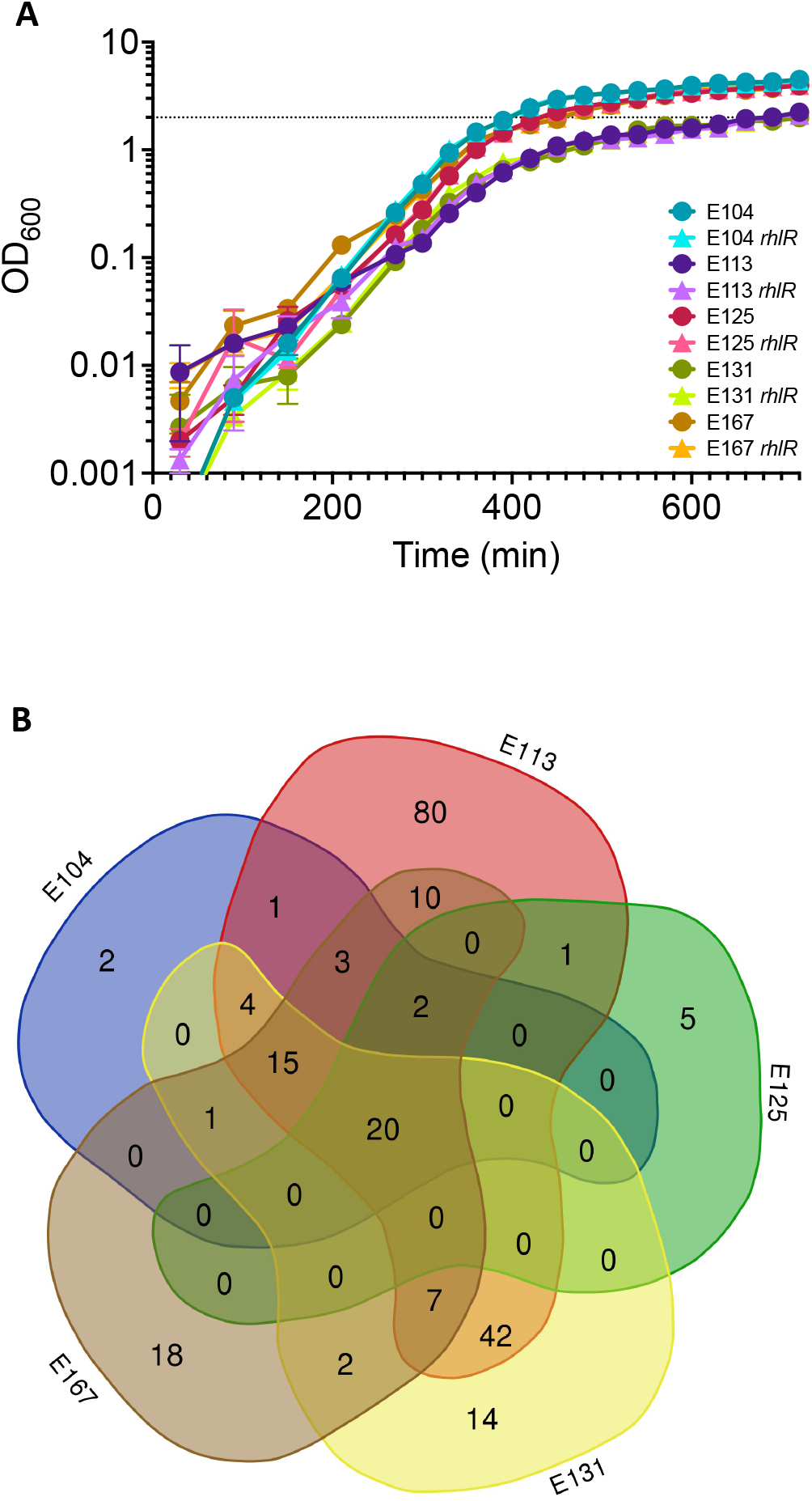
RhlR activated genes: core analysis. (A) Growth of CF isolate strains is similar between isogenic wild-type and *rhlR* mutant pairs. Strains were grown from a starting OD_600_ = 0.001 in LB+MOPS medium. Timing of total RNA harvest is shown with a dotted line (OD_600_ = 2.0). (B) Venn diagram displaying overlap of RhlR activated genes in individual CF isolate regulons. Lobes not scaled to size.

RhlR regulons varied in size from 28 (E125) to 185 genes (E113) (Table S4). A 5-way comparison revealed just 20 genes were shared among all 5 RhlR regulons, representing a core regulon for RhlR among our cohort of CF isolates (Figure 4B). This core RhlR regulon is detailed in Table 2. Genes in the core regulon include those coding for several known virulence factors that have been previously reported to be regulated by RhlR[19, 26]. These include genes coding for rhamnolipid (*rhlA*, PA3479), elastase (*lasB*, PA3724), hydrogen cyanide (*hcnABC*,PA2193-2195), and phenazine biosynthesis (*phzA1-B1*, PA4210-4211). *rhlI* (PA3476) was found to be under strong positive regulation by RhlR, indicating the autoinduction loop present in LasR/LasI in the Las system of other strains is also present in the Rhl system of our cohort isolates. The only gene in the core regulon annotated as a downstream transcriptional regulator is *vqsR* (PA2591). Comprising nearly half of the core regulon, the 8 RhlR-regulated genes spanning PA3326-3334 encode a recently described azetidomonamide (*aze*) biosynthesis regulon (PA3331 is RhlR-regulated in 4/5 isolates)[32, 33]. Finally, the core regulon also includes genes coding for two hypothetical proteins (PA1221 and PA4141). Expanding the core RhlR regulon list to those genes regulated by at least 4 of the 5 isolates (17 genes) adds the second phenazine locus (*phzA2B2C2*), LasA protease (*lasA*), and genes involved in T6SS (*hsiA2*) and rhamnolipid biosynthesis (*rhlB*). The average fold-change of the 20 core genes activated by RhlR was roughly 59-fold across the isolate cohort, but varied considerably by isolate, ranging from 4.93 in E125 to 219.28 in E113 (Table 2).

**Table 2.** Core RhlR regulated genes among CF isolates.

| Locus tag<sup>1</sup> | Gene name<sup>2</sup> | Function<sup>2</sup> | CF isolate RhIR-induced fold-change |  |  |  |  |
| --- | --- | --- | --- | --- | --- | --- | --- |
|  |  |  | E104 | E113 | E125 | E131 | E167 |
| PA1221 |  | hypothetical protein | 5.19 | 20.07 | 2.71 | 4.03 | 5.78 |
| PA1869 |  | probable acyl carrier protein | 28.72 | 79.34 | 5.78 | 14.77 | 10.05 |
| PA2193 | *hcnA* | hydrogen cyanide synthase HcnA | 23.85 | 65.55 | 9.66 | 8.91 | 39.16 |
| PA2194 | *hcnB* | hydrogen cyanide synthase HcnB | 8.19 | 8.61 | 2.35 | 2.92 | 6.93 |
| PA2195 | *hcnC* | hydrogen cyanide synthase HcnC | 3.43 | 8.52 | 2.06 | 2.74 | 5.53 |
| PA2591 | *vqsR* | virulence and quorum sensing regulator VqsR | 13.63 | 10.87 | 6.89 | 3.34 | 14.16 |
| PA3326 | *azeA/cIpP2* | AzeA/CIpP2 | 16.10 | 19.85 | 2.18 | 2.85 | 13.51 |
| PA3327 | *azeB* | probable non-ribosomal peptide synthetase | 21.13 | 102.17 | 2.83 | 5.12 | 36.02 |
| PA3328 | *azeC* | probable FAD-dependent monooxygenase | 67.86 | 271.93 | 5.28 | 12.05 | 152.08 |
| PA3329 | *azeD* | hypothetical protein | 18.71 | 83.98 | 3.03 | 8.22 | 45.88 |
| PA3330 | *azeE* | probable short chain dehydrogenase | 30.19 | 148.95 | 4.06 | 8.44 | 73.99 |
| PA3332 | *azeG* | conserved hypothetical protein | 38.93 | 98.62 | 3.96 | 18.39 | 70.46 |
| PA3333 | *azeH/fabH2* | AzeH 3-oxoacyl-[acyl-carrier-protein] synthase III | 22.65 | 73.33 | 2.15 | 14.31 | 40.57 |
| PA3334 |  | probable acyl carrier protein | 20.35 | 103.48 | 3.10 | 17.32 | 36.68 |
| PA3476 | *rhlI* | autoinducer synthesis protein RhII | 58.66 | 33.05 | 19.30 | 17.59 | 47.91 |
| PA3479 | *rhlA* | rhamnosyltransferase chain A | 18.19 | 84.86 | 6.79 | 20.95 | 13.62 |
| PA3724 | *lasB* | elastase LasB | 11.07 | 103.15 | 3.31 | 25.49 | 15.78 |
| PA4141 |  | hypothetical protein | 3.80 | 23.59 | 2.69 | 63.25 | 4.67 |
| PA4210 | *phzA1* | probable phenazine biosynthesis protein | 8.34 | 1788.72 | 6.27 | 13.02 | 63.05 |
| PA4211 | *phzB1* | probable phenazine biosynthesis protein | 23.71 | 1256.93 | 4.25 | 39.82 | 50.65 |

<sup>1</sup>PAO1 locus tags as determined by sequence homology.
<sup>2</sup>Gene names and functions from Pseudomonas.com, with aze annotation from Patteson et al. 2019.

In addition to the positively regulated genes described above, we sought to determine if any commonalities existed between the genes found to be repressed by RhlR in each isolate. RhlR-repressed regulons varied in size from 1 gene (E125) to 43 genes (E113), and in combination yielded a total of 69 genes across all 5 regulons. However, no genes were shared among all 5 regulons and only a single gene was found to be shared among any two strains (Figure S3), consistent with the idea that RhlR is a transcriptional activator[34].

To better understand the scope of RhlR regulation in our cohort of CF isolates, we combined all 5 CF isolate RhlR regulons to create a RhlR panregulon of 227 genes (Table S4). This RhlR panregulon list includes 26 genes not currently annotated in the PAO1 genome; 14 genes are not present in the PAO1 genome, while the remaining 12 are present in the PAO1 genome but not annotated as open reading frames (ORFs). Six of these acquired genes are positively regulated by RhlR in more than one isolate. There are also 3 putative operons controlled by RhlR in this acquired gene list: a 2 gene operon in E125 (287.9771.peg.292-293), a separate two gene operon in E131 (2913690790-1) and a 6 gene operon in E113 (2913679565-70).

Our observations of variability in the scope of QS regulation among our CF isolate cohort led us to question what genes might represent direct RhlR regulation. We found 30 of the 201 genes in the RhlR panregulon (28%, genes annotated in PAO1 only) harbor a putative binding site for RhlR upstream of the coding sequence, and an additional 33 genes are in predicted operons with a putative RhlR binding site in the promoter upstream of the first gene (Table S4). Of these 63, 18 directly RhlR-regulated genes are in the RhlR core regulon (90% of core regulon).

## Discussion

Infection of the CF airways by *P. aeruginosa* presents an intractable clinical problem. A growing body of evidence suggests that adaptation by *P. aeruginosa* to the CF infection niche includes a rewiring of the QS gene regulatory network observed in laboratory strains, with deleterious mutation of *lasR* leaving RhlR as the pivotal QS regulator. Unlike *lasR*, mutation of RhlR in CF isolates is uncommon[15, 27], which implies an evolutionary pressure for maintaining a functional Rhl-QS system as isolates transition to a more chronic phenotype. Thus, an understanding of mechanisms of RhlR liberation from LasR control, as well as the conserved targets of RhlR regulation, present an inflection point in efforts to control *P. aeruginosa* infections.

We focused our genetic analysis on a group of *lasR*-null CF isolates in an attempt to understand how QS might be rewired in lung infections. In our pangenomic analysis, we did not find an obvious link between the common genes lost or gained relative to laboratory strains that could explain this rewiring. Indeed, no known modulator of QS was found to be mutated in all our isolates except *lasR* itself. However, many candidates for this function remain. Mutation of the drug-efflux regulator *mexT* was previously shown to facilitate RhlR independence in *lasR-*null PAO1, although the precise mechanism that leads to increased RhlR activation in that background is still mysterious[28, 29]. However, like E90 in our initial description of Las-independent RhlR QS, the isolates in this study harbor functional *mexT* alleles[26]. Other regulators of multidrug efflux were found to be mutated across our isolate cohort (*armR, oprR, cmrA*), and it is possible that a combination of mutations other than in *mexT* could yield a mechanistically similar effect.

More broadly, our analysis provides a framework to approach future studies of genome evolution in CF-adapted *P. aeruginosa*. Our analysis includes isolates from a pediatric study that followed young individuals with CF after acquisition of *P. aeruginosa* infection. Genomic analysis indicated that adaptation is rapid in these populations; all 5 CF isolates in the current study harbored deleterious mutations in 197 genes, many coding for acute virulence factors. In addition to *lasR*, mutation of genes coding for O-antigen biosynthesis, exotoxin A, type III secretion system components, multidrug efflux, and twitching motility was present in our CF isolates when compared with the laboratory strain PAO1. Mutation of genes in each of these categories was previously observed during the adaptation of a clonal lineage of infecting *P. aeruginosa* studied in a pediatric patient, followed from 12 months to 96 months of age[15]. We also found common mutations in genes coding for S-pyocin production, a bacteriocin mechanism that allows *P. aeruginosa* to toxify neighboring non-kin strains using proteinaceous antimicrobial peptides internalized via siderophore receptors such as FptA[35, 36]. Divergence of genes coding for S-pyocins may reflect the pressures of interbacterial competition within infections, as selection may also target other similar competitive mechanisms such as the phage-tail-like R-pyocins or T4SS in evolving infection isolates[37]. The regional isolation of specific lineages of *P. aeruginosa* found in a study of explanted CF lungs provides additional support for this view of isolate pathoadaptation; as bacteria become isolated to a specific longterm niche in the lung, selection may purify mechanisms associated with competitive exclusion[38].

Unlike descriptions in laboratory strains[39], *lasR* mutation does not necessarily yield QS-null phenotypes in *P. aeruginosa[40]*. Here, we show inactivating mutations in *lasR* can be present in multiple strains that still activate QS target promoters through RhlR and can subsequently produce strong QS phenotypes. Mutation of *lasR* has been previously connected to the early stages of chronic CF infections, and the timing of collection of the EPIC study isolates used here agree with that notion [15, 41, 42]. Studies that focused on trends in *P. aeruginosa* virulence phenotypes associated with worsening lung disease have generally found these phenotypes to be attenuated. Pyocyanin and protease production were associated more with new onset or intermittent infection isolates than chronic isolates in an epidemiological survey that was part of the EPIC study, and a related study associated *lasR* mutation with both these phenotypes and with failure to eradicate *P. aeruginosa*[43, 44]. Our isolate cohort exhibited nominal changes in protease activity and significantly increased pyocyanin in every isolate, in contrast to the general trends observed in those earlier studies. Las-independent RhlR control of such phenotypes does not explain this discrepancy, but our data provide an alternate framework for understanding *lasR* mutation in long-lived infections. What previously was assumed to be an evolutionary path of QS inactivation may instead be a retuning of the QS network in chronic infection.

Our transcriptome analysis exploited these CF *P. aeruginosa* isolates to understand and generalize the scope of RhlR gene regulation. We found 20 genes encoding the production of a minimum of 7 products in common among isolates in our RhlR core regulon analysis. In addition to gene promoters known to be tightly controlled by RhlR (*rhlA*, rhamnolipid biosynthesis), our results confirmed RhlR control of many genes previously described as key virulence factors controlled by Las-dominated QS; *lasB* elastase, hydrogen cyanide biosynthesis (*hcnABC*), and phenazine biosynthesis (*phzA1B1*) are now understood to be regulated at least in part by RhlR. Our results also firmly position the azetidomonamide biosynthesis locus (*aze*, PA3326-34) under transcriptional control by RhlR. Indeed, nearly half of the core RhlR regulon is dedicated to these biosynthetic genes. While the primary biosynthetic product of this non-ribosomal-peptide-synthetase (NRPS) containing cluster has recently been described to be azabicyclene, among other congeners, the biological importance of this novel group of compounds to *P. aeruginosa* is unknown [32, 33]. Presence of the gene coding for the C4-HSL signal synthase RhlI in the core regulon provides evidence for a Rhl-QS circuit capable of autoregulatory feedback, an established feature of the LasR-I system[45–47]. We are interested in discerning the features of the RhlR-I autoregulatory loop in future investigations of Las-independent QS.

The gene coding for the transcriptional regulator VqsR was the only such annotated gene discovered in our core RhlR regulon analysis. VqsR has been previously found to be activated directly by LasR[48], and was also shown to provide a general activating effect on the QS circuitry through increased signal production[49]. However, a focused study showed this effect is indirect; VqsR does not directly bind the promoters of any recognized QS circuitry beyond the orphan regulator QscR[50]. Although VqsR was also found to directly bind the promoters of a small group of additional genes in that study, that VqsR would repress *qscR* to upregulate the QS circuitry fits with existing evidence surrounding such an anti-activation role for QscR[51–53]. Our results expand the control of *vqsR* beyond LasR to also include RhlR, and further demonstrate its conservation as a QS target in pathoadapted isolates, but additional studies will be necessary to disentangle the role of this downstream transcriptional activator in the interconnected QS networks of *P. aeruginosa*.

We noted several differences in RhlR regulon size and strength of regulation among our tested isolates, suggesting the conditions of RhlR QS control are diverse among strains with defective LasR. Strain E113 exhibited the greatest breadth and strength of RhlR control in our transcriptome analysis, with 185 genes activated and an average of 219-fold activation of the core RhlR regulon genes. Pyocyanin production in this strain was 25-fold greater than PAO1, and essentially absent upon deletion of *rhlR*, consistent with tight regulation of the *phzA1* locus in our transcriptome analysis (greater than 1000-fold). Yet the much smaller RhlR-activated foldchange value of approximately 15-fold exhibited in strain E131 was associated with similar pyocyanin phenotypes. This disparity suggests transcript fold-change values may be more indicative of tightness of regulation than reflective of phenotypes.

We were interested in how our RhlR core regulon analysis compared to two previous studies of QS gene regulation in CF isolates. We first compared the core RhlR regulon to that of our previously published analysis of the RhlR regulon of isolate E90 [26]. E90 was shown to positively regulate 53 genes in that analysis, and addition of that regulon to our core analysis did not change the 20 core genes regulated by RhlR. We then compared the RhlR core regulon to the QS regulons reported by Chugani and colleagues for PAO1 and 4 environmental isolates (“Chugani environmental core”, 5 strains, 42 genes) and a core QS regulon among 2 CF isolates (“Chugani CF core”, 2 strains, 25 genes) [31]. The Chugani environmental core shares 12 genes with our RhlR core regulon, including *hcnABC, rhlI, lasB, vqsR*, and much of the *aze* biosynthetic cluster (Figure 5A). In contrast, the Chugani CF core shares just 5 genes with our RhlR core regulon, including *lasB, vqsR*, and 3 genes of the *aze* operon (Figure 5B).

**Figure 5.**
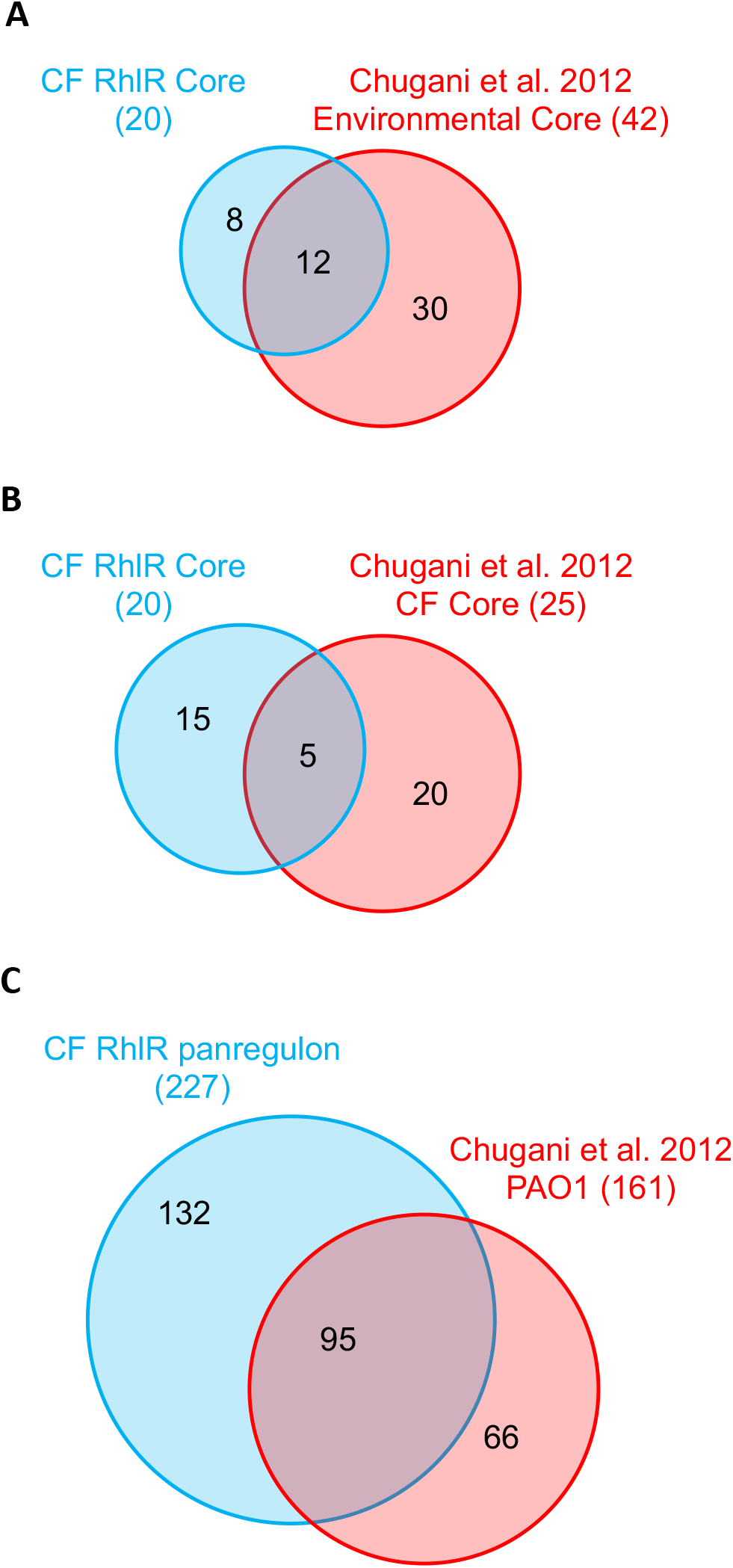
RhlR regulon comparisons. Comparison of the CF isolate RhlR core regulon determined here with (A) a core QS regulon of 4 environmental strains and PAO1 and (B) a core QS regulon of 2 CF isolates. (C) Comparison of the CF isolate RhlR panregulon with a PAO1 QS regulon. Comparison regulons from Chugani et al. 2012. Venn diagram lobes in each panel are scaled to approximate proportions.

We combined all 5 CF isolate RhlR regulons to create a RhlR panregulon of 227 genes, providing a more complete scope of RhlR regulation in our isolate cohort (Table S4). Twenty-six of these genes are not present in PAO1, indicating the scope of QS regulatory targets is more broad than that identified in laboratory strains. We then compared the remaining 201 genes of the RhlR panregulon to the PAO1 QS transcriptome described in the study by Chugani and colleagues, as that study used comparable conditions in determination of that reference QS regulon[31]. In that study, PAO1 was found to positively regulate 161 genes, 95 of which (59%) are represented in our RhlR panregulon (Figure 5C, Table S4). The remaining 106 RhlR-regulated genes that are annotated in PAO1, but not reported as differentially regulated by QS by Chugani and colleagues, are overrepresented for metabolic resources related to niche adaptation. These include genes involved in anaerobic and microaerophilic growth (*ccoN2*, *ccoO2*, *nrdD*), ethanol oxidation (*eraS*), nitrous oxide (*nosY*) and nitrate (*nirJ*) reduction, and components of the oxidative stress response (*fprA, soxR*). A principal goal of our study was to discern stable, heritable commonalities in RhlR gene regulation among CF isolates as opposed to condition-specific regulation, which guided our transcriptomics approach of standard laboratory media and conditions. That such niche specific adaptations are clearly present in the RhlR regulons of our tested isolates in laboratory conditions suggests a robust but adaptable relationship between QS and infections. We also queried the relationship between our RhlR panregulon genes present in PAO1 and a transcriptomic microarray study that studied multiple sampling timepoints and multiple configurations of QS-on or -off conditions by Schuster and colleagues [19]. Similar to comparison with the Chugani study, we found 62% of our RhlR-regulated genes (125/201) present in PAO1 were also upregulated in at least one condition at least one timepoint in that study. Our panregulon also revealed RhlR controls the gene coding for the quinolone monooxygenase PqsH in multiple isolates, an enzyme previously reported to be under LasR, but not RhlR control in laboratory strains[12]. PqsH oxidizes the relatively low-affinity PqsR ligand 2-heptyl-4-hydroxyquinoline (HHQ) to produce the relatively high-affinity ligand 2-heptyl-3-hydroxy-4(1H)-quinolone (PQS)[54], providing a potential avenue for RhlR to tune PQS signalling in some CF isolates. Together, our panregulon analyses suggest that while timing of QS regulation may account for some differences in regulon content, adaptation to a specific environment may also have lasting effects on the scope of QS regulation.

We conclude that clear commonalities exist among Las-independent RhlR QS in CF-adapted *P. aeruginosa*. Twenty genes encoding 7 products, including potent virulence factors, were regulated by RhlR in our analysis of 5 CF isolates exhibiting this circuitry. Our results position RhlR as an independent QS regulator of virulence genes in *P. aeruginosa*, reaffirming the potential for this QS receptor as a therapeutic target[55].

## Materials and methods

### Isolate selection, growth, and characterization

Bacterial strains and plasmids used in this study are listed in Table S5. The CF infection isolates were originally collected from oropharyngeal and sputum samples from patients 5 to 12 years of age as part of the Early *Pseudomonas* Infection Control (EPIC) observational study[21, 43]. Routine cultures were maintained on Luria-Bertani (LB) agar medium at 37°C or in LB broth buffered with 50 mM 3-(*N-* morpholino)propanesulfonic acid (MOPS) at pH 6.8 (LB+MOPS) in 18 mm borosilicate glass culture tubes with 250 rpm orbital shaking at 37°C, unless otherwise noted. When appropriate, gentamicin was added at 10 μg/ml for *Escherichia coli* (Gm10) or 100 μg/ml for *P. aeruginosa* (Gm100). For synthetic signal addition experiments, 2 μM *N*-3-oxo-dodecanoyl-L-homoserine lactone (3OC12-HSL) or 10 μM *N*-butanoyl-L-homoserine lactone (C4-HSL) (Cayman Chemical) were added.

### Transcriptional fusions and phenotypic analysis

LasR and RhlR transcriptional activity was determined using P*_lasI_-gfp* or P*_rhlA_-gfp* transcriptional fusion reporter plasmids, respectively, as described previously[17]. Plasmids were introduced into electrocompetent *P. aeruginosa* cells prepared using repeated 300 mM sucrose washes, selected on LB Gm100 agar, and verified by PCR[56]. For endpoint experiments, individual gentamicin-resistant colonies were inoculated into LB+MOPS broth with or without synthetic signal and grown at 37°C with 250 rpm shaking, and then harvested at 18 h. 200 μl triplicate aliquots were then transferred to black-walled chimney-welled fluorescence microtiter plates with optically clear bottoms (Greiner Bio-one) and assayed in a BioTek Synergy HI microplate reader for optical density at 600 nm (OD_600_) and GFP-derived fluorescence (GFP; λ_excitation_=485 nm, λ_emission_=535 nm). For kinetic experiments, from overnight cultures of isolated colonies in LB+MOPS Gm100, cells were subcultured 1:100 into fresh LB+MOPS Gm100 and grown to mid logarithmic phase (OD_600_ of 0.2-0.8, 1 cm pathlength), and then inoculated into 200 μl LB+MOPS broth with or without synthetic signal at a starting OD_600_ of 0.01 in triplicate in fluorescence microtiter plates. Plates were then incubated with orbital shaking at 37°C in a plate reader with OD_600_ and GFP measurements every 15 min for 16 h. Reported fluorescence was normalized to OD_600_, and fluorescence derived from each strain with a promoterless control reporter plasmid of the same backbone was subtracted to account for background fluorescence. All experiments were performed with a minimum of biological triplicates.

Pyocyanin production was determined after 18 h of growth with shaking at 37°C in pyocyanin production medium (PPM) as previously described[17]. Extracellular protease production was determined by patching isolated colonies onto skim milk agar plates (1/4 strength LB broth/4% wt/vol skim milk/1.5%wt/vol agar, 20 ml in 150 mm petri dishes)[24], followed by incubation at 37°C for 48 h, and then the zone of proteolytic clearing measured using a ruler from the edge of the colony. A minimum of 3 biological replicates were performed, and laboratory strain controls were included on every plate.

### *rhlR* mutant construction

Markerless in-frame *rhlR* deletion mutants were made using homologous recombination-mediated allelic exchange as previously described[57, 58]. Strain-specific *rhlR* deletion plasmids were constructed using *E. coli*-mediated assembly of PCR-amplified products[59]. Initial native *rhlR* sequences were determined using PCR-amplification and targeted Sanger sequencing (Azenta-Genewiz) of 4 overlapping fragments covering a ~2.5 kb region encompassing the *rhlR* locus, which were then used to design sequence-specific deletion allele construction primers. Allele fragments were generated using the high-fidelity PrimeSTAR Max polymerase (Takara Bio), assembled into strain-specific pEXG2 plasmids in *E. coli* DH5α (New England Biolabs), and introduced into each isolate using *sacB-* mediated sucrose counterselection as previously described[58, 59]. Deletion was confirmed using PCR amplification and targeted sequencing.

### Genomic analysis

High-molecular weight (HMW) genomic DNA was isolated from liquid cultures using the Qiagen Genomic-tip 20/G kit (Qiagen). DNA was then subjected to a hybrid approach of both short- and long-read DNA sequencing to achieve high-confidence complete *de novo* genome assemblies, as we have reported previously[26]. Short reads (Illumina MiSeq, TruSeq v3, 300-bp paired-end; Illumina) were groomed using FastQC (Babraham Bioinformatics, Babraham Institute) in combination with Trimmomatic (v0.36; adapter trimming, paired reads only, Phred score cutoff = 15)[60]. After basecalling and demultiplexing using Guppy (v3.1.5, Oxford Nanopore), long reads (EXP-PBC001, EXP-NBD114 libraries, Nanopore MinION, R9.4.1 pores, Oxford

Nanopore) were groomed using NanoPack (NanoPlot v1.27.0; NanoQC v0.9.1) and Porechop (v0.2.4)[61, 62]. Hybrid-assembly mode in the Unicycler pipeline (including SPAdes v3.13.0, Racon v1.4.3, Pilon) was used to assemble and close genomes prior to annotation using the RAST pipeline[63–66]. General genomic features are summarized in Table S1.

Comparative genomic analysis was achieved using a combination of established tools. Pangenomic analysis was conducted using complete genomes in the program “anvi-pan-genome” in the anvi’o software ecosystem[67]. PAO1 genes not present in all isolates (74) were discovered using the “anvi-display-pan” program. Variant analysis was conducted using Illumina short reads compared against reference strains (PAO1, accession NC_002516; PA14, accession GCF_000014625.1) using the StrandNGS SNP, CNV, and SV pipelines (v3.3.1, Strand Life Sciences). Variants were defined with a variant read frequency >90% and indels defined as <100 bp. We defined variants as present in cohort isolates but not laboratory strains by first comparing each genome to PAO1, followed by Venn analysis to determine a core group of variants (1682), followed by filtering against PA14 variants to yield a final list of 123 genes.

We conducted a phylogenetic analysis using the comparative genomics suite available through the Joint Genome Institute Integrated Microbial Genomes and Microbiomes – Expert Review (JGI IMG/MER) webtool analysis suite. The phylogeny was constructed using hierarchical clustering in the Taxonomy-Genus mode, and the presence/absence of GC-1 and −2 in genomes was determined using the genomic Conserved Neighborhood analysis tool.

### Transcriptomic analysis

We determined the extent of QS-controlled gene regulation in our cohort of 5 CF isolates that exhibit Las-independent RhlR QS activity using an RNA-seq-based transcriptome analysis. We compared engineered *rhlR* deletion strains (QS-off condition) to isogenic parent strains supplemented with both synthetic C4-HSL (10 μM) and 3OC12-HSL (2 μM) to ensure QS activation (QS-on). We did so because, while isolates in our cohort do not respond to 3OC12-HSL through LasR (Figure 2), the orphan LuxR homolog QscR is still intact and likely binds to exogenous 3OC12-HSL from neighboring bacteria in the context of infections. QscR has been shown to bind and activate a single linked operon beginning with PA1897[51], and its role in RhlR QS regulation in these isolates is unknown.

Beginning with isolated colonies, we grew each strain overnight in 3 ml LB+MOPS in 18 mm culture tubes at 37°C with 250 rpm shaking. Saturated cultures were then diluted to OD_600_=0.001 in 25 ml LB+MOPS medium (with AHL signal addition in QS-on samples) in 250 ml aerobic baffled flasks and incubated at 37°C with 250 rpm shaking. Total RNA was harvested from approximately 1 × 10^9^ cells in the transition from logarithmic to stationary phase (OD_600_=2.0). Total RNA was preserved and purified using RNAprotect Bacteria Reagent and an RNeasy Mini Kit with Qiazol (Qiagen) as previously described[31]. RNA was quantified using Qubit fluorometric quantitation (Thermo Fisher Scientific) and lack of RNA degradation confirmed using NorthernMax-Gly glyoxal gel electrophoresis (Thermo Fisher Scientific) and an Agilent 2100 Bioanalyzer (Agilent). rRNA depletion, library generation, and >20 M 150-bp paired-end Illumina HiSeq reads were generated for each sample commercially (Azenta-Genewiz). Reads were groomed using FastQC (Babraham Bioinformatics, Babraham Institute) and Trim Galore! (v0.4.3; https://github.com/FelixKrueger/TrimGalore). We mapped reads from 3 biological replicates to the strain-specific mapping references generated earlier using the Subread/featureCounts suite of command line tools, and conducted differential expression (DE) analyses in the R statistical environment using strain-specific datasets in the DESeq2 statistical package with a log2-foldchange cutoff of 1 and a false discovery rate (FDR) of 0.05[68–70]. We used an all-by-all blastp comparison to determine the common gene names associated with targets in PAO1, and where possible we use those names here[71]. Genes with blastp top hit bitscores less than 50 were considered non-homologous to any PAO1 gene. *rhlR* was discarded from analysis post-computation of DE, as this is a result of our engineered knockouts in the QS-off condition. *qscR* and the single linked target of QscR, the operon spanning PA1897-91, were also discarded from analysis as differential regulation of these genes is due to synthetic 3OC12-HSL addition in the QS-on condition, as discussed above. Venn analysis and core figure generation was conducted using the online tool provided by Bioinformatics & Evolutionary Genomics, University of Gent, Belgium (https://bioinformatics.psb.ugent.be/webtools/Venn/).

### Promoter analysis

A global position analysis has not yet been published for RhlR, but a consensus promoter sequence for binding of LuxR homologs has been determined to have dyad symmetry of N2CTN12AGN2 centered 38-45 bp upstream of the transcriptional start site (TSS)[72, 73]. LasR binding sites have been explored in a global position analysis using chromatin immunoprecipitation combined with DNA microarrays (ChIP-chip) and through direct *in vitro* binding in electrophoretic mobility shift assays [74]. LasR and RhlR share LuxR homology and have been shown to regulate many of the same promoters, so we assumed a *las* or *lux*-box recognition site may also be directly regulated by RhlR (“*las/rhl*-box”). We cross-referenced our RhlR panregulon with a previous study that mapped the transcription start sites of *P. aeruginosa* strain PA14 to determine whether a gene is predicted to lie in a solely transcribed ORF or if the gene is the first in an operon[75]. We then searched the PRODORIC, CollecTF, and PseudoCAP databases to determine if a RhlR-regulated gene, or the first gene in an operon containing a RhlR-regulated gene, has a putative *las/rhl-box* [76].

## Data availability

All genomic (RefSeq) and transcriptomic (GEO) data associated with this study are publicly available through NCBI under BioProject PRJNA784987. CF isolate genomes are also available through the Integrated Microbial Genomes/Expert Review database (Joint Genome Institute) under the following genome IDs: E104, 2913679597; E113, 2913673089; E125, 2870682680; E131, 2913685979; E167, 2870695665.

## Acknowledgements

This work was supported by US Public Health Service (USPHS) Grant R01GM125714 (to A.A.D.). A.A.D. was also supported by the Burroughs-Wellcome Fund (Grant 1012253) and Doris Duke Charitable Foundation (Grant 2017073). K.L.A. was supported by a Postdoctoral Fellowship from the Cystic Fibrosis Foundation (ASFAHL19F0). We acknowledge core support from USPHS Grant P30DK089507 and the Cystic Fibrosis Foundation (Grants SINGH15R0 and R565 CR11).

## Notes

### Competing Interest Statement

The authors have declared no competing interest.

