## Supplemental tables and figures for "Genetic and transcriptomic characteristics of RhlR-dependent quorum sensing in cystic fibrosis isolates of *Pseudomonas aeruginosa*"

**Table S1. CF isolate genomic features.**

| Strain | Length (bp) | Features[1] | Plasmid (bp) |
|---|---|---|---|
| PAO1 | 6264403 | 5901 | - |
| E104 | 6680448 | 6414 | - |
| E113 | 6864839 | 6544 | - |
| E125 | 6895810 | 6798 | - |
| E131 | 6666693 | 6379 | - |
| E167 | 6831875 | 6661 | 47445 |

[1]Features enumerated from RAST annotations.

**Table S2. PAO1 genes with nonsynonymous mutations in all CF isolates.**

| Locus tag[1] | Gene name[2] | Function[2] | Full deletion |
|---|---|---|---|
| PA0078 | *tssL1* | TssL1 | |
| PA0084 | *tssC1* | TssC1 | |
| PA0093 | *tse6* | Tse6 | |
| PA0170 | | hypothetical protein | |
| PA0225 | | probable transcriptional regulator | |
| PA0241 | | probable major facilitator superfamily (MFS) transporter | |
| PA0256 | | hypothetical protein | |
| PA0285 | | conserved hypothetical protein | |
| PA0287 | *gpuP* | 3-guanidinopropionate transport protein | |
| PA0373 | *ftsY* | signal recognition particle receptor FtsY | |
| PA0457 | | hypothetical protein | |
| PA0548 | *tktA* | transketolase | |
| PA0558 | | conserved hypothetical protein | |
| PA0559 | | conserved hypothetical protein | |
| PA0561 | | hypothetical protein | Yes |
| PA0715 | | hypothetical protein | Yes |
| PA0716 | | hypothetical protein | Yes |
| PA0717 | | hypothetical protein of bacteriophage Pf1 | Yes |
| PA0724 | | probable coat protein A of bacteriophage Pf1 | Yes |
| PA0729 | | hypothetical protein | Yes |
| PA0787 | | hypothetical protein | |
| PA0807 | *ampDh3* | AmpDh3 | |
| PA0842 | | probable glycosyl transferase | |
| PA0860 | | probable ATP-binding/permease fusion ABC transporter | Yes |
| PA0919 | | alanyl-phosphatidylglycerol hydrolase | Yes |
| PA0946 | | hypothetical protein | Yes |
| PA0948 | | hypothetical protein | |
| PA0977 | | hypothetical protein | Yes |
| PA0983 | | conserved hypothetical protein | Yes |
| PA0985 | *pyoS5* | pyocin S5 | Yes |
| PA1012 | | conserved hypothetical protein | |
| PA1014 | *wapB* | 1,2-glucosyltransferase WapB | |
| PA1019 | *mucK* | cis,cis-muconate transporter MucK | |
| PA1128 | | probable transcriptional regulator | |
| PA1150 | *pys2* | pyocin S2 | |
| PA1218 | | hypothetical protein | |
| PA1219 | | hypothetical protein | |
| PA1229 | | probable transcriptional regulator | |
| PA1240 | | probable enoyl-CoA hydratase/isomerase | |
| PA1245 | *aprX* | AprX | |
| PA1283 | | probable transcriptional regulator | |
| PA1286 | | probable major facilitator superfamily (MFS) transporter | |
| PA1356 | | hypothetical protein | |
| PA1368 | | hypothetical protein | Yes |
| PA1369 | | hypothetical protein | Yes |
| PA1370 | | hypothetical protein | Yes |
| PA1371 | | hypothetical protein | Yes |
| PA1372 | | hypothetical protein | Yes |
| PA1381 | | hypothetical protein | Yes |
| PA1382 | *xqhB* | probable type II secretion system protein | Yes |
| PA1383 | | hypothetical protein | Yes |
| PA1384 | *galE* | UDP-glucose 4-epimerase | Yes |
| PA1385 | | probable glycosyl transferase | Yes |

| | | | |
|---|---|---|---|
| PA1386 | | probable ATP-binding component of ABC transporter | Yes |
| PA1387 | | hypothetical protein | Yes |
| PA1388 | | hypothetical protein | Yes |
| PA1389 | | probable glycosyl transferase | Yes |
| PA1390 | | probable glycosyl transferase | Yes |
| PA1391 | | probable glycosyl transferase | Yes |
| PA1392 | | hypothetical protein | Yes |
| PA1393 | *cysC* | adenosine 5'-phosphosulfate (APS) kinase | Yes |
| PA1416 | | conserved hypothetical protein | |
| PA1430 | *lasR* | transcriptional regulator LasR | |
| PA1495 | | hypothetical protein | |
| PA1614 | *gpsA* | glycerol-3-phosphate dehydrogenase, biosynthetic | |
| PA1620 | | hypothetical protein | |
| PA1621 | | probable hydrolase | |
| PA1714 | *exsD* | ExsD | Yes |
| PA1721 | *pscH* | type III export protein PscH | |
| PA1749 | | hypothetical protein | |
| PA1813 | | probable hydroxyacylglutathione hydrolase | |
| PA1842 | | hypothetical protein | |
| PA1887 | | hypothetical protein | Yes |
| PA1888 | | hypothetical protein | Yes |
| PA1892 | | hypothetical protein | |
| PA1909 | | hypothetical protein | |
| PA1914 | | conserved hypothetical protein | |
| PA1935 | | hypothetical protein | Yes |
| PA1939 | | hypothetical protein | Yes |
| PA2047 | *cmrA* | probable transcriptional regulator | |
| PA2048 | | hypothetical protein | |
| PA2073 | | probable transporter (membrane subunit) | Yes |
| PA2074 | | hypothetical protein | Yes |
| PA2118a | | hypothetical protein | |
| PA2139 | | hypothetical protein | Yes |
| PA2192 | | conserved hypothetical protein | |
| PA2218 | | hypothetical protein | Yes |
| PA2219 | *opdE* | membrane protein OpdE | Yes |
| PA2220 | *oprR* | probable transcriptional regulator | Yes |
| PA2229 | | conserved hypothetical protein | |
| PA2233 | *pslC* | PslC | |
| PA2335 | | probable TonB-dependent receptor | |
| PA2336 | | hypothetical protein | |
| PA2340 | | probable binding-protein-dependent maltose/mannitol transport prot | Yes |
| PA2403 | *fpvG* | FpvG | |
| PA2497 | | probable transcriptional regulator | |
| PA2618 | | hypothetical protein | |
| PA2626 | *trmU* | tRNA methyltransferase | |
| PA2707 | | hypothetical protein | |
| PA2730 | | hypothetical protein | Yes |
| PA2731 | | Uncharacterized protein | Yes |
| PA2732 | | hypothetical protein | Yes |
| PA2733 | | conserved hypothetical protein | Yes |
| PA2734 | | hypothetical protein | Yes |
| PA2735 | | probable restriction-modification system protein | Yes |
| PA2736 | | hypothetical protein | Yes |
| PA2745 | | probable hydrolase | |
| PA2813 | | probable glutathione S-transferase | |

| | | | |
|---|---|---|---|
| PA2846 | | probable transcriptional regulator | |
| PA2870 | | diguanylate cyclase | |
| PA2885 | *atuR* | putative repressor of atu genes | |
| PA2910 | | conserved hypothetical protein | |
| PA2922 | | probable hydrolase | |
| PA3146 | *wbpK* | probable NAD-dependent epimerase/dehydratase WbpK | Yes |
| PA3147 | *wbpJ* | probable glycosyl transferase WbpJ | Yes |
| PA3148 | *wbpI* | UDP-N-acetylglucosamine 2-epimerase WbpI | Yes |
| PA3149 | *wbpH* | probable glycosyltransferase WbpH | Yes |
| PA3150 | *wbpG* | LPS biosynthesis protein WbpG | Yes |
| PA3151 | *hisF2* | imidazoleglycerol-phosphate synthase, cyclase subunit | Yes |
| PA3152 | *hisH2* | glutamine amidotransferase | Yes |
| PA3153 | *wzx* | O-antigen translocase | Yes |
| PA3154 | *wzy* | B-band O-antigen polymerase | Yes |
| PA3157 | | probable acetyltransferase | Yes |
| PA3163 | *cmk* | cytidylate kinase | |
| PA3187 | | probable ATP-binding component of ABC transporter | |
| PA3285 | | probable sigma-70 factor, ECF subfamily | |
| PA3302 | | conserved hypothetical protein | |
| PA3364 | *amiC* | aliphatic amidase expression-regulating protein | |
| PA3373 | | conserved hypothetical protein | |
| PA3374 | | conserved hypothetical protein | |
| PA3407 | *hasAp* | heme acquisition protein HasAp | |
| PA3455 | | conserved hypothetical protein | |
| PA3492 | | conserved hypothetical protein | |
| PA3565 | | probable transcriptional regulator | Yes |
| PA3589 | | probable acyl-CoA thiolase | |
| PA3598 | | conserved hypothetical protein | |
| PA3670 | | hypothetical protein | |
| PA3719 | *armR* | antirepressor for MexR, ArmR | Yes |
| PA3780 | | hypothetical protein | Yes |
| PA3790 | *oprC* | Putative copper transport outer membrane porin OprC precursor | |
| PA3840 | | conserved hypothetical protein | |
| PA3843 | | hypothetical protein | Yes |
| PA3866 | | pyocin S4 | Yes |
| PA3867 | | probable DNA invertase | Yes |
| PA3868 | | hypothetical protein | Yes |
| PA3869 | | hypothetical protein | Yes |
| PA3890 | *opuCB* | OpuC ABC transporter, permease protein, OpuCB | |
| PA3893 | | conserved hypothetical protein | |
| PA3931 | | conserved hypothetical protein | |
| PA3932 | | probable transcriptional regulator | Yes |
| PA3933 | *betT3* | BetT3 | |
| PA3959 | | hypothetical protein | |
| PA4014 | | hypothetical protein | |
| PA4188 | | conserved hypothetical protein | |
| PA4192 | | probable ATP-binding component of ABC transporter | |
| PA4193 | | probable permease of ABC transporter | |
| PA4195 | | probable binding protein component of ABC transporter | |
| PA4211 | *phzB1* | probable phenazine biosynthesis protein | |
| PA4313a | | hypothetical protein | |
| PA4330 | | probable enoyl-CoA hydratase/isomerase | |
| PA4442 | *cysN* | ATP sulfurylase GTP-binding subunit/APS kinase | |
| PA4466 | | probable phosphoryl carrier protein | |
| PA4525 | *pilA* | type 4 fimbrial precursor PilA | Yes |

| | | | |
|---|---|---|---|
| PA4526 | *pilB* | type 4 fimbrial biogenesis protein PilB | |
| PA4546 | *pilS* | two-component sensor PilS | |
| PA4549 | *fimT* | type 4 fimbrial biogenesis protein FimT | |
| PA4554 | *pilY1* | type 4 fimbrial biogenesis protein PilY1 | |
| PA4620 | | hypothetical protein | |
| PA4628 | *lysP* | lysine-specific permease | |
| PA4642 | | hypothetical protein | |
| PA4664 | *prmC* | S-adenosylmethionine-dependent methyltransferase, PrmC | |
| PA4714 | | conserved hypothetical protein | |
| PA4730 | *panC* | pantoate--beta-alanine ligase | |
| PA4790 | | conserved hypothetical protein | |
| PA4797 | | probable transposase | Yes |
| PA4872 | | hypothetical protein | |
| PA4884 | | hypothetical protein | |
| PA4911 | | probable permease of ABC branched-chain amino acid transporter | |
| PA4914 | *amaR* | transcriptional regulator, AmaR | |
| PA4916 | *nrtR* | Nudix-related transcriptional regulator NrtR | |
| PA4988 | *waaA* | 3-deoxy-D-manno-octulosonic-acid (KDO) transferase | |
| PA5030 | | probable major facilitator superfamily (MFS) transporter | |
| PA5069 | *tatB* | translocation protein TatB | |
| PA5082 | *dguC* | DguC | |
| PA5085 | *dguR* | DguR | |
| PA5112 | *estA* | esterase EstA | |
| PA5122 | | hypothetical protein | |
| PA5141 | *hisA* | phosphoribosylformimino-5-aminoimidazole carboxamide | Yes |
| PA5246 | | conserved hypothetical protein | |
| PA5263 | *argH* | argininosuccinate lyase | Yes |
| PA5296 | *rep* | ATP-dependent DNA helicase Rep | Yes |
| PA5345 | *recG* | ATP-dependent DNA helicase RecG | |
| PA5349 | *rubB* | rubredoxin reductase | |
| PA5382 | | probable transcriptional regulator | Yes |
| PA5418 | *soxA* | sarcosine oxidase alpha subunit | |
| PA5500 | *znuC* | zinc transport protein ZnuC | |
| PA5502 | | hypothetical protein | |

[1]PAO1 locus tags as determined by sequence homology.

[2]Gene names and functions from Pseudomonas.com.

**Table S3. Genes present in all isolates, but not present in PAO1.**

| Strain-specific gene ID[a] | | | | | | | |
|---|---|---|---|---|---|---|---|
| E104 | E113 | E125 | E131 | E167 | Gene length (bp) | Gene cluster | Annotated function[a] |
| 2913680624 | 2913676237 | 2870687275 | 2913687009 | 2870696693 | 100 | | hypothetical protein |
| 2913680625 | 2913676417 | 2870685150 | 2913687010 | 2870696694 | 98 | | transposase |
| 2913680836 | 2913674339 | 2870683918 | 2913687221 | 2870696905 | 402 | | Integrase |
| 2913681549 | 2913675136 | 2870684674 | 2913687875 | 2870697559 | 319 | | nucleoside-diphosphate-sugar epimerase |
| 2913679820 | 2913673343 | 2870682899 | 2913686204 | 2870695889 | 145 | | hypothetical protein |
| 2913682201 | 2913676275 | 2870685342 | 2913688528 | 2870697914 | 1039 | | Cu(I)/Ag(I) efflux system membrane protein CusA/SilA |
| 2913682210 | 2913676267 | 2870685351 | 2913688537 | 2870697923 | 691 | | Cu+-exporting ATPase |
| 2913682302 | 2913677723 | 2870685390 | 2913688582 | 2870697968 | 464 | | integrating conjugative element protein (TIGR03755 family) |
| 2913682303 | 2913677722 | 2870685391 | 2913688583 | 2870697969 | 316 | | integrating conjugative element protein (TIGR03756 family) |
| 2913682326 | 2913677702 | 2870685414 | 2913688606 | 2870697992 | 730 | | conjugative coupling factor TraD (TOL family) |
| 2913682337 | 2913677684 | 2870685420 | 2913688617 | 2870698003 | 759 | | SNF2 family DNA or RNA helicase |
| 2913682748 | 2913676288 | 2870685965 | 2913689046 | 2870698819 | 180 | GC-1 | type VI protein secretion system component VasK |
| 2913682749 | 2913676289 | 2870685966 | 2913689047 | 2870698820 | 399 | GC-1 | DNA-binding PucR family transcriptional regulato |
| 2913682750 | 2913676290 | 2870685967 | 2913689048 | 2870698821 | 288 | GC-1 | ectoine hydroxylase-related dioxygenase (phytanoyl-CoA dioxygenase family) |
| 2913682751 | 2913676291 | 2870685968 | 2913689049 | 2870698822 | 320 | GC-1 | photosystem II stability/assembly factor-like uncharacterized protein |
| 2913682752 | 2913676292 | 2870685969 | 2913689050 | 2870698823 | 800 | GC-1 | predicted RND superfamily exporter protein |
| 2913682753 | 2913676293 | 2870685970 | 2913689051 | 2870698824 | 548 | GC-1 | hypothetical protein |
| 2913682754 | 2913676294 | 2870685971 | 2913689052 | 2870698825 | 474 | GC-1 | hypothetical protein |
| 2913682755 | 2913676295 | 2870685972 | 2913689053 | 2870698826 | 220 | GC-1 | hypothetical protein |
| 2913682756 | 2913676296 | 2870685973 | 2913689054 | 2870698827 | 455 | GC-1 | glycine/D-amino acid oxidase-like deaminating enzyme |
| 2913682757 | 2913676297 | 2870685974 | 2913689055 | 2870698828 | 295 | GC-1 | ectoine hydroxylase-related dioxygenase (phytanoyl-CoA dioxygenase family |
| 2913682758 | 2913676298 | 2870685975 | 2913689056 | 2870698829 | 193 | GC-1 | AcrR family transcriptional regulator |
| 2913682759 | 2913676299 | 2870685976 | 2913689057 | 2870698830 | 508 | GC-1 | cation diffusion facilitator CzcD-associated flavoprotein CzcO |
| 2913682760 | 2913676300 | 2870685977 | 2913689058 | 2870698831 | 291 | GC-1 | pimeloyl-ACP methyl ester carboxylesterase |
| 2913682761 | 2913676301 | 2870685978 | 2913689059 | 2870698832 | 385 | GC-1 | alcohol dehydrogenase |
| 2913682762 | 2913676302 | 2870685979 | 2913689060 | 2870698833 | 443 | GC-1 | amino acid transporter |
| 2913682763 | 2913676303 | 2870685980 | 2913689061 | 2870698834 | 474 | GC-1 | aminobutyraldehyde dehydrogenase |
| 2913682764 | 2913676304 | 2870685981 | 2913689062 | 2870698835 | 417 | GC-1 | diaminobutyrate-2-oxoglutarate transaminase |
| 2913682765 | 2913676305 | 2870685982 | 2913689063 | 2870698836 | 319 | GC-1 | 3-hydroxyisobutyrate dehydrogenase-like beta-hydroxyacid dehydrogenase |
| 2913682766 | 2913676306 | 2870685983 | 2913689064 | 2870698837 | 50 | GC-1 | hypothetical protein |
| 2913682767 | 2913676307 | 2870685984 | 2913689065 | 2870698838 | 552 | GC-1 | cation/acetate symporter |
| 2913682768 | 2913676308 | 2870685985 | 2913689066 | 2870698839 | 107 | GC-1 | uncharacterized membrane protein (DUF485 family) |
| 2913682769 | 2913676309 | 2870685986 | 2913689067 | 2870698840 | 549 | GC-1 | fatty-acyl-CoA synthase |
| 2913682770 | 2913676310 | 2870685987 | 2913689068 | 2870698841 | 255 | GC-1 | NAD(P)-dependent dehydrogenase (short-subunit alcohol dehydrogenase family |
| 2913682771 | 2913676311 | 2870685988 | 2913689069 | 2870698842 | 390 | GC-1 | acyl-CoA dehydrogenase |
| 2913682772 | 2913676312 | 2870685989 | 2913689070 | 2870698843 | 118 | GC-1 | hypothetical protein |
| 2913682773 | 2913676313 | 2870685990 | 2913689071 | 2870698844 | 351 | GC-1 | aminoglycoside phosphotransferase (APT) family kinase protein |
| 2913682774 | 2913676314 | 2870685991 | 2913689072 | 2870698845 | 650 | GC-1 | propionate catabolism operon transcriptional regulator |
| 2913682775 | 2913676315 | 2870685992 | 2913689073 | 2870698846 | 167 | GC-1 | hypothetical protein |
| 2913682864 | 2913676395 | 2870686081 | 2913689161 | 2870698935 | 71 | | hypothetical protein |
| 2913683112 | 2913676672 | 2870686330 | 2913689409 | 2870699183 | 291 | | 4-hydroxy-tetrahydrodipicolinate synthase |
| 2913683662 | 2913677091 | 2870686877 | 2913689959 | 2870699822 | 50 | | hypothetical protein |
| 2913684055 | 2913677628 | 2870688001 | 2913690352 | 2870700216 | 288 | GC-2 | chromosome partitioning related protein ParA |
| 2913684056 | 2913677629 | 2870688002 | 2913690353 | 2870700217 | 245 | GC-2 | hypothetical protein |
| 2913684057 | 2913677630 | 2870688003 | 2913690354 | 2870700218 | 165 | GC-2 | hypothetical protein |
| 2913684058 | 2913677631 | 2870688004 | 2913690355 | 2870700219 | 239 | GC-2 | hypothetical protein |
| 2913684060 | 2913677632 | 2870688006 | 2913690357 | 2870700221 | 233 | GC-2 | hypothetical protein |
| 2913684061 | 2913677635 | 2870688009 | 2913690358 | 2870700222 | 465 | GC-2 | replicative DNA helicase |
| 2913684062 | 2913677636 | 2870688011 | 2913690359 | 2870700223 | 61 | GC-2 | formate dehydrogenase maturation protein FdhE |
| 2913684063 | 2913677637 | 2870688012 | 2913690360 | 2870700224 | 175 | GC-2 | hypothetical protein |

| | | | | | | | |
|---|---|---|---|---|---|---|---|
| 2913684064 | 2913677638 | 2870688013 | 2913690361 | 2870700225 | 85 | GC-2 | hypothetical protein |
| 2913684065 | 2913677639 | 2870688014 | 2913690362 | 2870700226 | 79 | GC-2 | hypothetical protein |
| 2913684066 | 2913677640 | 2870688015 | 2913690363 | 2870700227 | 77 | GC-2 | hypothetical protein |
| 2913684068 | 2913677643 | 2870688017 | 2913690365 | 2870700229 | 89 | GC-2 | hypothetical protein |
| 2913684069 | 2913677644 | 2870688018 | 2913690366 | 2870700230 | 576 | GC-2 | ParB family protein of integrating conjugative element (PFGI_1 class) |
| 2913684070 | 2913677645 | 2870688019 | 2913690367 | 2870700231 | 255 | GC-2 | hypothetical protein |
| 2913684071 | 2913677646 | 2870688020 | 2913690368 | 2870700232 | 441 | GC-2 | hypothetical protein |
| 2913684075 | 2913677647 | 2870688023 | 2913690372 | 2870700236 | 242 | GC-2 | integrating conjugative element protein (TIGR03761 family) |
| 2913684076 | 2913677648 | 2870688024 | 2913690373 | 2870700237 | 177 | GC-2 | hypothetical protein |
| 2913684077 | 2913677650 | 2870688026 | 2913690374 | 2870700238 | 162 | GC-2 | single-strand DNA-binding protein |
| 2913684079 | 2913677651 | 2870688033 | 2913690376 | 2870700240 | 639 | GC-2 | DNA topoisomerase-1 |
| 2913684081 | 2913677654 | 2870688035 | 2913690378 | 2870700242 | 629 | GC-2 | hypothetical protein |
| 2913684082 | 2913677655 | 2870688036 | 2913690379 | 2870700243 | 657 | GC-2 | hypothetical protein |
| 2913684083 | 2913677656 | 2870688039 | 2913690380 | 2870700244 | 374 | GC-2 | type IV pili sensor histidine kinase/response regulator |
| 2913684084 | 2913677657 | 2870688040 | 2913690381 | 2870700245 | 569 | GC-2 | type IVB pilus formation R64 PilN family outer membrane protein |
| 2913684085 | 2913677658 | 2870688041 | 2913690382 | 2870700246 | 441 | GC-2 | hypothetical protein |
| 2913684086 | 2913677659 | 2870688042 | 2913690383 | 2870700247 | 177 | GC-2 | type IV pilus biogenesis protein PilP |
| 2913684087 | 2913677660 | 2870688043 | 2913690384 | 2870700248 | 526 | GC-2 | type II secretory ATPase GspE/PulE/Tfp pilus assembly ATPase PilB-like protein |
| 2913684088 | 2913677661 | 2870688044 | 2913690385 | 2870700249 | 359 | GC-2 | type II secretory pathway component PulF |
| 2913684089 | 2913677662 | 2870688045 | 2913690386 | 2870700250 | 176 | GC-2 | type II secretory pathway pseudopilin PulG |
| 2913684090 | 2913677663 | 2870688046 | 2913690387 | 2870700251 | 313 | GC-2 | twitching motility protein PilT |
| 2913684091 | 2913677664 | 2870688047 | 2913690388 | 2870700252 | 442 | GC-2 | type II secretory pathway pseudopilin PulG |
| 2913684092 | 2913677665 | 2870688048 | 2913690389 | 2870700253 | 145 | GC-2 | hypothetical protein |
| 2913684094 | 2913677667 | 2870688050 | 2913690391 | 2870700255 | 129 | GC-2 | hypothetical protein |
| 2913684095 | 2913677668 | 2870688051 | 2913690392 | 2870700256 | 58 | GC-2 | hypothetical protein |
| 2913684098 | 2913677671 | 2870688054 | 2913690395 | 2870700259 | 162 | GC-2 | hypothetical protein |
| 2913684100 | 2913677673 | 2870688056 | 2913690397 | 2870700261 | 65 | GC-2 | hypothetical protein |
| 2913684102 | 2913677675 | 2870688057 | 2913690399 | 2870700263 | 235 | GC-2 | hypothetical protein |
| 2913684104 | 2913677677 | 2870688059 | 2913690401 | 2870700265 | 116 | GC-2 | hypothetical protein |
| 2913684105 | 2913677678 | 2870688060 | 2913690402 | 2870700266 | 256 | GC-2 | hypothetical protein |
| 2913684106 | 2913677680 | 2870688061 | 2913690403 | 2870700267 | 120 | GC-2 | hypothetical protein |
| 2913684107 | 2913677681 | 2870688062 | 2913690404 | 2870700268 | 84 | GC-2 | hypothetical protein |
| 2913684108 | 2913677682 | 2870688063 | 2913690405 | 2870700269 | 201 | GC-2 | hypothetical protein |
| 2913684109 | 2913677683 | 2870688064 | 2913690406 | 2870700270 | 395 | GC-2 | hypothetical protein |
| 2913684117 | 2913676282 | 2870685438 | 2913690414 | 2870700278 | 116 | GC-2 | mercuric ion transport protein |
| 2913684135 | 2913677696 | 2870688073 | 2913690432 | 2870700296 | 90 | GC-2 | hypothetical protein |
| 2913684136 | 2913677697 | 2870688074 | 2913690433 | 2870700297 | 229 | GC-2 | soluble cytochrome b562 |
| 2913684137 | 2913677698 | 2870688075 | 2913690434 | 2870700298 | 251 | GC-2 | integrating conjugative element protein (TIGR03759 family |
| 2913684138 | 2913677699 | 2870688076 | 2913690435 | 2870700299 | 193 | GC-2 | hypothetical protein |
| 2913684139 | 2913677700 | 2870688077 | 2913690436 | 2870700300 | 166 | GC-2 | integrating conjugative element protein (TIGR03765 family) |
| 2913684140 | 2913677701 | 2870688078 | 2913690437 | 2870700301 | 89 | GC-2 | hypothetical protein |
| 2913684142 | 2913677703 | 2870688080 | 2913690439 | 2870700303 | 248 | GC-2 | integrating conjugative element membrane protein (TIGR03747 family) |
| 2913684143 | 2913677705 | 2870688081 | 2913690440 | 2870700304 | 493 | GC-2 | superfamily I DNA/RNA helicase |
| 2913684144 | 2913677706 | 2870688082 | 2913690441 | 2870700305 | 369 | GC-2 | 3-dehydroquinate dehydratase |
| 2913684145 | 2913677707 | 2870688083 | 2913690442 | 2870700306 | 99 | GC-2 | hypothetical protein |
| 2913684146 | 2913677708 | 2870688084 | 2913690443 | 2870700307 | 97 | GC-2 | RAQPRD family integrative conjugative element protein |
| 2913684147 | 2913677709 | 2870688085 | 2913690444 | 2870700308 | 49 | GC-2 | integrating conjugative element protein (TIGR03758 family) |
| 2913684148 | 2913677710 | 2870688086 | 2913690445 | 2870700309 | 73 | GC-2 | integrating conjugative element membrane protein (TIGR03745 family) |
| 2913684149 | 2913677711 | 2870688087 | 2913690446 | 2870700310 | 128 | GC-2 | conjugative transfer region protein (TIGR03750 family) |
| 2913684150 | 2913677712 | 2870688088 | 2913690447 | 2870700311 | 219 | GC-2 | integrating conjugative element protein (TIGR03746 family) |
| 2913684151 | 2913677713 | 2870688089 | 2913690448 | 2870700312 | 294 | GC-2 | integrating conjugative element protein (TIGR03749 family) |
| 2913684152 | 2913677714 | 2870688090 | 2913690449 | 2870700313 | 501 | GC-2 | integrating conjugative element protein (TIGR03752 family) |
| 2913684153 | 2913677715 | 2870688091 | 2913690450 | 2870700314 | 147 | GC-2 | conjugative transfer region lipoprotein (TIGR03751 family) |

| | | | | | | | |
|---|---|---|---|---|---|---|---|
| 2913684154 | 2913677716 | 2870688092 | 2913690451 | 2870700315 | 980 | GC-2 | conjugative transfer ATPase |
| 2913684155 | 2913677717 | 2870688093 | 2913690452 | 2870700316 | 94 | GC-2 | hypothetical protein |
| 2913684159 | 2913677721 | 2870688099 | 2913690456 | 2870700320 | 63 | GC-2 | integrating conjugative element protein (TIGR03757 family) |
| 2913684162 | 2913677724 | 2870687290 | 2913690459 | 2870700323 | 115 | GC-2 | Kef-type K+ transport system membrane component KefB |
| 2913684163 | 2913677725 | 2870687291 | 2913690460 | 2870700324 | 511 | GC-2 | hypothetical protein |
| 2913684166 | 2913677727 | 2870687293 | 2913690463 | 2870700327 | 90 | GC-2 | antitoxin ParD1/3/4 |
| 2913684167 | 2913677728 | 2870687294 | 2913690464 | 2870700328 | 116 | GC-2 | toxin ParE1/3/4 |
| 2913684173 | 2913677734 | 2870687295 | 2913690470 | 2870700334 | 639 | GC-2 | integrating conjugative element relaxase (TIGR03760 family) |
| 2913684174 | 2913677735 | 2870687296 | 2913690471 | 2870700335 | 426 | GC-2 | Integrase |

[a]Gene IDs and annotated functions from the IMG/MER database.

**Table S4. RhlR panregulon.**

| | Locus Tag[a] | Gene name[a] | Product Name[a] | Fold-change[b] | | | | | | | | | |
|---|---|---|---|---|---|---|---|---|---|---|---|---|---|
| | | | | E104 | | E113 | | E125 | | E131 | | 167 | |
| | | | | FC | FCSE | FC | FCSE | FC | FCSE | FC | FCSE | FC | FCSE |
| | PA0050 | | hypothetical protein | | | 3.36 | 1.23 | | | 2.39 | 1.27 | | |
| | PA0051 | *phzH* | potential phenazine-modifying enzyme | | | 3.22 | 1.24 | | | 4.25 | 1.21 | | |
| • | PA0052 | | hypothetical protein | | | 2.01 | 1.22 | | | 2.99 | 1.17 | | |
| | PA0111 | | hypothetical protein | | | | | | | | | 2.42 | 1.24 |
| • ○ | PA0122 | *rahU* | rahU | 5.03 | 1.14 | 48.53 | 1.33 | | | 26.13 | 1.33 | 6.47 | 1.11 |
| | PA0123 | | probable transcriptional regulator | | | 2.80 | 1.18 | | | | | | |
| | PA0130 | *bauC* | 3-Oxopropanoate dehydrogenase | | | 2.18 | 1.24 | | | | | | |
| | PA0132 | *bauA* | Beta-alanine:pyruvate transaminase | | | 2.71 | 1.31 | | | | | | |
| | PA0195 | *pntAA* | putative NAD(P) transhydrogenase, subunit alpha part 1 | | | 2.04 | 1.17 | | | | | | |
| | PA0195.1 | *pntAB* | putative NAD(P) transhydrogenase, subunit alpha part 2 | | | 2.56 | 1.17 | | | | | | |
| | PA0197 | *tonB2* | TonB2 | | | | | 2.19 | 1.23 | | | | |
| | PA0198 | *exbB1* | transport protein ExbB | | | 2.31 | 1.21 | | | | | | |
| | PA0199 | *exbD1* | transport protein ExbD | | | 3.34 | 1.22 | | | | | | |
| | PA0200 | | hypothetical protein | | | 4.80 | 1.24 | | | | | | |
| | PA0208 | *mdcA* | malonate decarboxylase alpha subunit | | | 3.95 | 1.35 | | | | | | |
| | PA0209 | | conserved hypothetical protein | | | 4.11 | 1.48 | | | | | | |
| | PA0210 | *mdcC* | malonate decarboxylase delta subunit | | | 6.97 | 1.46 | | | | | | |
| | PA0211 | *mdcD* | malonate decarboxylase beta subunit | | | 6.31 | 1.79 | | | | | | |
| | PA0213 | | hypothetical protein | | | 3.46 | 1.46 | | | | | | |
| | PA0214 | | probable acyl transferase | | | 2.78 | 1.32 | | | | | | |
| | PA0511 | *nirJ* | heme d1 biosynthesis protein NirJ | | | | | | | | | 2.63 | 1.23 |
| | PA0534 | *pauB1* | FAD-dependent oxidoreductase | | | | | | | | | 2.81 | 1.15 |
| | PA0546 | *metK* | methionine adenosyltransferase | | | 2.04 | 1.17 | | | | | | |
| | PA0547 | | probable transcriptional regulator | | | 2.46 | 1.11 | | | | | | |
| • | PA0852 | *cbpD* | chitin-binding protein CbpD precursor | 2.01 | 1.12 | 8.73 | 1.24 | | | 8.93 | 1.11 | 2.52 | 1.11 |
| | PA0865 | *hpd* | 4-hydroxyphenylpyruvate dioxygenase | | | | | | | | | 5.02 | 1.56 |
| | PA0866 | *aroP2* | aromatic amino acid transport protein AroP2 | | | | | | | | | 4.71 | 1.23 |
| | PA0870 | *phhC* | aromatic amino acid aminotransferase | | | | | | | | | 2.83 | 1.36 |
| | PA0871 | *phhB* | pterin-4-alpha-carbinolamine dehydratase | | | | | | | | | 3.19 | 1.35 |
| • ○ | PA0997 | *pqsB* | PqsB | | | | | 2.31 | 1.12 | | | | |
| • ○ | PA0998 | *pqsC* | PqsC | | | | | | | | | 2.04 | 1.25 |
| ○ | PA0999 | *pqsD* | 3-oxoacyl-[acyl-carrier-protein] synthase III | | | | | | | | | 2.22 | 1.19 |
| • ○ | PA1000 | *pqsE* | Quinolone signal response protein | | | | | | | | | 2.64 | 1.13 |
| • ○ | PA1130 | *rhlC* | rhamnosyltransferase 2 | | | 2.85 | 1.16 | | | 2.47 | 1.13 | | |
| • ○ | PA1131 | | probable major facilitator superfamily (MFS) transporter | | | 3.72 | 1.20 | | | | | | |
| | PA1168 | | hypothetical protein | | | 5.01 | 1.44 | | | | | | |
| | PA1212 | | probable major facilitator superfamily (MFS) transporter | | | 4.31 | 1.16 | | | 2.53 | 1.25 | | |
| | PA1213 | | hypothetical protein | | | 5.97 | 1.17 | | | 3.32 | 1.34 | | |
| | PA1214 | | hypothetical protein | 2.40 | 1.29 | 12.32 | 1.23 | | | 4.15 | 1.30 | | |
| | PA1215 | | hypothetical protein | 5.23 | 1.16 | 8.83 | 1.15 | | | 4.73 | 1.32 | | |
| • | PA1216 | | hypothetical protein | 7.75 | 1.14 | 36.49 | 1.19 | | | 12.30 | 1.47 | 2.91 | 1.13 |
| • | PA1217 | | probable 2-isopropylmalate synthase | | | 3.32 | 1.15 | | | | | | |
| • | PA1218 | | hypothetical protein | | | 10.85 | 1.15 | | | 4.50 | 1.36 | | |
| | PA1219 | | hypothetical protein | | | 18.84 | 1.21 | | | 3.72 | 1.32 | 2.46 | 1.32 |
| | PA1220 | | hypothetical protein | 3.23 | 1.30 | 19.64 | 1.19 | | | 3.61 | 1.33 | 3.45 | 1.24 |
| • | **PA1221** | | **hypothetical protein** | **5.19** | **1.20** | **20.07** | **1.16** | **2.71** | **1.19** | **4.03** | **1.27** | **5.78** | **1.14** |
| • ○ | PA1245 | *aprX* | AprX | 2.21 | 1.09 | 3.68 | 1.27 | | | 5.32 | 1.23 | | |
| • ○ | PA1246 | *aprD* | alkaline protease secretion protein AprD | | | 3.31 | 1.24 | | | 4.51 | 1.21 | | |
| • ○ | PA1247 | *aprE* | alkaline protease secretion protein AprE | | | 2.90 | 1.20 | | | 6.27 | 1.27 | | |
| • ○ | PA1248 | *aprF* | Alkaline protease secretion outer membrane protein AprF precursor | | | 2.36 | 1.23 | | | 4.18 | 1.25 | | |
| • | PA1249 | *aprA* | alkaline metalloproteinase precursor | | | 2.48 | 1.27 | | | 5.02 | 1.21 | | |
| • | PA1250 | *aprI* | alkaline proteinase inhibitor AprI | | | 2.23 | 1.19 | | | 3.31 | 1.23 | | |
| | PA1317 | *cyoA* | cytochrome o ubiquinol oxidase subunit II | | | 2.42 | 1.17 | | | | | | |
| | PA1319 | *cyoC* | cytochrome o ubiquinol oxidase subunit III | | | 2.18 | 1.24 | | | | | | |
| • ○ | PA1431 | *rsaL* | regulatory protein RsaL | | | | | | | 2.52 | 1.26 | | |
| • ○ | PA1432 | *lasI* | autoinducer synthesis protein LasI | | | | | | | 2.28 | 1.19 | 2.52 | 1.15 |
| | PA1546 | *hemN* | oxygen-independent coproporphyrinogen III oxidase | | | 2.31 | 1.27 | | | | | | |
| | PA1550 | | hypothetical protein | | | 2.64 | 1.17 | | | | | | |
| | PA1551 | | probable ferredoxin | | | 2.47 | 1.12 | | | | | | |
| | PA1556 | *ccoO2* | Cytochrome c oxidase, cbb3-type, CcoO subunit | | | 2.59 | 1.26 | | | | | | |

| | | | | | | | | | | | | | | |
|---|---|---|---|---|---|---|---|---|---|---|---|---|---|---|
| | | PA1557 | *ccoN2* | Cytochrome c oxidase, cbb3-type, CcoN subunit | | | 3.25 | 1.15 | | | | | | |
| • | ○ | PA1656 | *hsiA2* | HsiA2 | 4.73 | 1.10 | 18.47 | 1.15 | 2.05 | 1.12 | | | 4.27 | 1.13 |
| • | ○ | PA1657 | *hsiB2* | HsiB2 | 2.57 | 1.15 | 9.74 | 1.15 | | | | | 4.51 | 1.22 |
| • | ○ | PA1658 | *hsiC2* | HsiC2 | | | 4.83 | 1.19 | | | | | 2.56 | 1.13 |
| | ○ | PA1659 | *hsiF2* | HsiF2 | 2.33 | 1.25 | 5.22 | 1.20 | | | | | 2.69 | 1.19 |
| • | ○ | PA1660 | *hsiG2* | HsiG2 | | | 3.42 | 1.13 | | | | | | |
| • | ○ | PA1661 | *hsiH2* | HsiH2 | | | 4.80 | 1.16 | | | | | | |
| | ○ | PA1662 | *clpV2* | clpV2 | | | 2.93 | 1.12 | | | | | | |
| • | ○ | PA1663 | *sfa2* | Sfa2 | | | 4.36 | 1.16 | | | | | 2.09 | 1.21 |
| | ○ | PA1664 | *orfX* | OrfX | | | 5.32 | 1.20 | | | 4.91 | 1.60 | | |
| • | ○ | PA1665 | *fha2* | Fha2 | | | 5.15 | 1.19 | | | | | 2.26 | 1.17 |
| | ○ | PA1666 | *lip2* | Lip2 | | | 3.60 | 1.23 | | | | | | |
| | ○ | PA1668 | *dotU2* | DotU2 | | | 2.21 | 1.14 | | | | | | |
| | ○ | PA1670 | *stp1* | Stp1 | | | 2.38 | 1.19 | | | | | 2.04 | 1.15 |
| • | | PA1784 | | hypothetical protein | | | 2.04 | 1.27 | | | 2.00 | 1.14 | | |
| • | ○ | **PA1869** | | **probable acyl carrier protein** | **28.72** | **1.14** | **79.34** | **1.20** | **5.78** | **1.13** | **14.77** | **1.21** | **10.05** | **1.16** |
| | | PA1870 | | hypothetical protein | | | 2.15 | 1.21 | | | | | | |
| • | | PA1871 | *lasA* | LasA protease precursor | 2.48 | 1.13 | 13.94 | 1.21 | | | 5.85 | 1.14 | 2.25 | 1.10 |
| • | | PA1899 | *phzA2* | probable phenazine biosynthesis protein | 6.15 | 1.25 | 101.96 | 1.30 | | | 9.14 | 1.19 | 4.63 | 1.26 |
| • | | PA1900 | *phzB2* | probable phenazine biosynthesis protein | 4.60 | 1.18 | 49.57 | 1.31 | | | 12.33 | 1.16 | 2.64 | 1.15 |
| | | PA1901 | *phzC2* | phenazine biosynthesis protein PhzC | 4.62 | 1.38 | 130.13 | 1.20 | | | 9.22 | 1.23 | 2.46 | 1.21 |
| | | PA1902 | *phzD2* | phenazine biosynthesis protein PhzD | | | | | | | 13.88 | 1.26 | | |
| | | PA1903 | *phzE2* | phenazine biosynthesis protein PhzE | | | 26.88 | 2.57 | | | | | 3.30 | 1.35 |
| | | PA1904 | *phzF2* | probable phenazine biosynthesis protein | | | 12.09 | 1.33 | | | | | | |
| | | PA1905 | *phzG2* | probable pyridoxamine 5'-phosphate oxidase | | | 12.07 | 1.33 | | | 12.25 | 1.19 | | |
| | | PA1920 | *nrdD* | class III (anaerobic) ribonucleoside-triphosphate reductase subunit, NrdD | | | 2.74 | 1.26 | | | | | | |
| | | PA1979 | *eraS* | sensor kinase, EraS | | | 2.77 | 1.23 | | | | | | |
| • | | PA2030 | | hypothetical protein | | | | | | | | | 2.10 | 1.12 |
| | | PA2031 | | hypothetical protein | | | | | | | 2.14 | 1.20 | 2.29 | 1.12 |
| • | | PA2066 | | hypothetical protein | | | 2.34 | 1.24 | | | 3.92 | 1.25 | | |
| | | PA2067 | | probable hydrolase | | | 3.52 | 1.24 | | | 5.91 | 1.29 | | |
| • | | PA2068 | | probable major facilitator superfamily (MFS) transporter | | | 2.47 | 1.21 | | | 4.81 | 1.20 | | |
| • | | PA2069 | | probable carbamoyl transferase | | | 12.90 | 1.31 | | | 8.42 | 1.14 | | |
| • | | PA2146 | | conserved hypothetical protein | | | 2.26 | 1.29 | | | 4.14 | 1.38 | | |
| • | | PA2159 | | conserved hypothetical protein | | | | | | | 2.13 | 1.25 | | |
| • | ○ | **PA2193** | ***hcnA*** | **hydrogen cyanide synthase HcnA** | **23.85** | **1.29** | **65.55** | **1.22** | **9.66** | **1.25** | **8.91** | **1.30** | **39.16** | **1.28** |
| • | ○ | **PA2194** | ***hcnB*** | **hydrogen cyanide synthase HcnB** | **8.19** | **1.21** | **8.61** | **1.15** | **2.35** | **1.15** | **2.92** | **1.20** | **6.93** | **1.20** |
| • | ○ | **PA2195** | ***hcnC*** | **hydrogen cyanide synthase HcnC** | **3.43** | **1.13** | **8.52** | **1.13** | **2.06** | **1.12** | **2.74** | **1.22** | **5.53** | **1.10** |
| | | PA2273 | *soxR* | SoxR | | | 2.31 | 1.10 | | | | | | |
| • | | PA2274 | | hypothetical protein | | | 34.03 | 1.26 | | | 12.96 | 1.17 | | |
| | | PA2275 | | probable alcohol dehydrogenase (Zn-dependent) | | | 8.87 | 1.27 | | | 3.91 | 1.19 | | |
| • | | PA2300 | *chiC* | chitinase | | | 8.04 | 1.25 | | | 9.08 | 1.19 | | |
| • | ○ | PA2302 | *ambE* | AmbE | | | 2.91 | 1.36 | | | 3.53 | 1.18 | | |
| • | ○ | PA2303 | *ambD* | AmbD | | | 4.37 | 1.36 | | | 4.15 | 1.17 | | |
| • | ○ | PA2304 | *ambC* | AmbC | | | 3.43 | 1.33 | | | 3.74 | 1.20 | | |
| • | ○ | PA2305 | *ambB* | AmbB | | | 2.88 | 1.27 | | | 4.65 | 1.20 | | |
| | | PA2321 | | gluconokinase | | | | | 2.04 | 1.20 | | | | |
| | | PA2327 | | probable permease of ABC transporter | | | 2.37 | 1.22 | | | | | | |
| • | | PA2328 | | hypothetical protein | | | 3.78 | 1.26 | | | | | | |
| • | | PA2329 | | probable ATP-binding component of ABC transporter | | | 3.74 | 1.25 | | | | | | |
| • | | PA2330 | | hypothetical protein | | | 3.28 | 1.21 | | | | | | |
| • | | PA2331 | | hypothetical protein | | | 5.94 | 1.25 | | | | | | |
| | | PA2507 | *catA* | catechol 1,2-dioxygenase | | | | | | | | | 5.89 | 1.46 |
| | | PA2508 | *catC* | muconolactone delta-isomerase | | | | | | | | | 5.21 | 1.42 |
| | | PA2509 | *catB* | muconate cycloisomerase I | | | | | | | | | 4.52 | 1.43 |
| • | | PA2564 | | hypothetical protein | | | | | | | 2.57 | 1.09 | | |
| | | PA2565 | | hypothetical protein | | | | | | | 2.22 | 1.14 | | |
| • | | PA2566 | | conserved hypothetical protein | | | | | | | 2.37 | 1.17 | | |
| • | ○ | PA2570 | *lecA* | LecA | | | 16.50 | 1.25 | | | 2.03 | 1.21 | | |
| • | ○ | PA2587 | *pqsH* | probable FAD-dependent monooxygenase | | | 4.39 | 1.29 | | | 2.72 | 1.12 | | |
| • | | PA2588 | | probable transcriptional regulator | | | 4.37 | 1.14 | | | 3.36 | 1.13 | | |
| • | | PA2589 | | hypothetical protein | | | 2.26 | 1.22 | | | | | | |
| | | PA2590 | | hypothetical protein | 2.03 | 1.15 | | | | | | | | |
| • | ○ | **PA2591** | ***vqsR*** | **VqsR** | **13.63** | **1.12** | **10.87** | **1.32** | **6.89** | **1.10** | **3.34** | **1.21** | **14.16** | **1.08** |

| | | | | | | | | | | | | | |
|---|---|---|---|---|---|---|---|---|---|---|---|---|---|
| • | PA2592 | | probable periplasmic spermidine/putrescine-binding protein | 4.76 | 1.11 | 10.74 | 1.18 | 2.28 | 1.10 | | | 3.82 | 1.06 |
| | PA2593 | *qteE* | quorum threshold expression element, QteE | 2.03 | 1.20 | 12.76 | 1.17 | | | | | 2.03 | 1.13 |
| | PA2594 | | conserved hypothetical protein | | | 2.81 | 1.14 | | | | | | |
| | PA2788 | | probable chemotaxis transducer | | | 2.44 | 1.18 | | | | | | |
| • | PA3022 | | hypothetical protein | | | 2.23 | 1.10 | | | | | | |
| • ○ | PA3104 | *xcpP* | secretion protein XcpP | | | 2.16 | 1.12 | | | | | | |
| | PA3318 | | hypothetical protein | | | | | | | 2.17 | 1.24 | | |
| • | PA3325 | | conserved hypothetical protein | 2.80 | 1.13 | 2.30 | 1.13 | | | | | | |
| • ○ | **PA3326** | ***azeA/clpP2*** | **ClpP2** | **16.10** | **1.09** | **19.85** | **1.22** | **2.18** | **1.15** | **2.85** | **1.19** | **13.51** | **1.07** |
| • ○ | **PA3327** | *azeB* | **probable non-ribosomal peptide synthetase** | **21.13** | **1.13** | **102.17** | **1.10** | **2.83** | **1.15** | **5.12** | **1.30** | **36.02** | **1.07** |
| ○ | **PA3328** | *azeC* | **probable FAD-dependent monooxygenase** | **67.86** | **1.25** | **271.93** | **1.21** | **5.28** | **1.25** | **12.05** | **1.43** | **152.08** | **1.16** |
| • ○ | **PA3329** | *azeD* | **hypothetical protein** | **18.71** | **1.17** | **83.98** | **1.19** | **3.03** | **1.26** | **8.22** | **1.44** | **45.88** | **1.18** |
| ○ | **PA3330** | *azeE* | **probable short chain dehydrogenase** | **30.19** | **1.24** | **148.95** | **1.25** | **4.06** | **1.26** | **8.44** | **1.47** | **73.99** | **1.26** |
| • ○ | PA3331 | | cytochrome P450 | 3.41 | 1.14 | 45.99 | 1.25 | | | 3.63 | 1.36 | 21.25 | 1.16 |
| • ○ | **PA3332** | *azeG* | **conserved hypothetical protein** | **38.93** | **1.26** | **98.62** | **1.33** | **3.96** | **1.37** | **18.39** | **1.54** | **70.46** | **1.27** |
| • ○ | **PA3333** | ***azeH/fabH2*** | **3-oxoacyl-[acyl-carrier-protein] synthase III** | **22.65** | **1.17** | **73.33** | **1.30** | **2.15** | **1.21** | **14.31** | **1.48** | **40.57** | **1.16** |
| ○ | **PA3334** | | **probable acyl carrier protein** | **20.35** | **1.16** | **103.48** | **1.35** | **3.10** | **1.20** | **17.32** | **1.88** | **36.68** | **1.13** |
| ○ | PA3335 | | hypothetical protein | 13.42 | 1.14 | 26.91 | 1.32 | | | 4.02 | 1.39 | 19.14 | 1.17 |
| ○ | PA3336 | | probable major facilitator superfamily (MFS) transporter | 4.74 | 1.19 | 11.19 | 1.31 | | | 2.85 | 1.36 | 7.96 | 1.30 |
| • ○ | PA3361 | *lecB* | fucose-binding lectin PA-IIL | | | 26.43 | 1.23 | | | 9.48 | 1.44 | 2.55 | 1.13 |
| | PA3395 | *nosY* | NosY protein | | | | | | | 2.24 | 1.22 | | |
| | PA3397 | *fprA* | FprA | | | 2.92 | 1.13 | | | | | | |
| | PA3415 | | probable dihydrolipoamide acetyltransferase | | | | | | | | | 2.17 | 1.23 |
| | PA3441 | | probable molybdopterin-binding protein | | | 2.06 | 1.20 | | | | | | |
| | PA3475 | *pheC* | cyclohexadienyl dehydratase precursor | | | 3.49 | 1.13 | | | 2.26 | 1.16 | | |
| • ○ | **PA3476** | ***rhlI*** | **autoinducer synthesis protein RhlI** | **58.66** | **1.09** | **33.05** | **1.69** | **19.30** | **1.10** | **17.59** | **1.22** | **47.91** | **1.10** |
| • ○ | PA3478 | *rhlB* | rhamnosyltransferase chain B | 2.18 | 1.11 | 10.40 | 1.23 | | | 5.03 | 1.16 | 2.26 | 1.16 |
| • ○ | **PA3479** | ***rhlA*** | **rhamnosyltransferase chain A** | **18.19** | **1.17** | **84.86** | **1.24** | **6.79** | **1.12** | **20.95** | **1.11** | **13.62** | **1.17** |
| | PA3519 | | hypothetical protein | | | 2.17 | 1.26 | | | | | | |
| • | PA3520 | | hypothetical protein | | | 4.24 | 1.19 | | | 2.34 | 1.16 | | |
| | PA3677 | *mexJ* | MexJ | | | 2.10 | 1.10 | | | | | | |
| | PA3718 | | probable major facilitator superfamily (MFS) transporter | | | 3.06 | 1.17 | | | | | | |
| | PA3719 | *armR* | antirepressor for MexR, ArmR | | | 2.28 | 1.23 | | | | | | |
| | PA3720 | | hypothetical protein | | | 2.05 | 1.24 | | | | | | |
| • ○ | **PA3724** | ***lasB*** | **elastase LasB** | **11.07** | **1.10** | **103.15** | **1.36** | **3.31** | **1.11** | **25.49** | **1.17** | **15.78** | **1.12** |
| | PA3734 | | hypothetical protein | | | 5.64 | 1.17 | | | 4.38 | 1.13 | | |
| | PA4067 | *oprG* | Outer membrane protein OprG precursor | | | 2.27 | 1.21 | | | | | | |
| • | PA4078 | | probable nonribosomal peptide synthetase | | | 2.10 | 1.16 | | | 2.51 | 1.18 | | |
| | PA4127 | *hpcG* | 2-oxo-hept-3-ene-1,7-dioate hydratase | | | | | | | | | 2.08 | 1.17 |
| • | PA4128 | | conserved hypothetical protein | | | 9.01 | 1.18 | | | | | 11.22 | 1.24 |
| • | PA4129 | | hypothetical protein | | | 57.11 | 1.13 | | | 4.46 | 1.36 | 29.28 | 1.20 |
| • | PA4130 | | probable sulfite or nitrite reductase | 2.17 | 1.11 | 38.90 | 1.10 | | | 4.92 | 1.16 | 16.16 | 1.14 |
| • | PA4131 | | probable iron-sulfur protein | 2.68 | 1.12 | 32.80 | 1.12 | | | 2.77 | 1.13 | 15.48 | 1.12 |
| • | PA4132 | | conserved hypothetical protein | | | 8.51 | 1.12 | | | | | 3.64 | 1.16 |
| • | PA4133 | | cytochrome c oxidase subunit (cbb3-type) | | | 44.37 | 1.16 | | | | | 6.15 | 1.15 |
| • | PA4134 | | hypothetical protein | | | 20.29 | 1.16 | | | | | 2.36 | 1.26 |
| | PA4136 | | probable major facilitator superfamily (MFS) transporter | | | 2.21 | 1.14 | | | | | | |
| • | **PA4141** | | **hypothetical protein** | **3.80** | **1.07** | **23.59** | **1.16** | **2.69** | **1.08** | **63.25** | **1.38** | **4.67** | **1.16** |
| • | PA4142 | | probable secretion protein | | | 5.19 | 1.17 | | | 2.15 | 1.12 | | |
| | PA4143 | | probable toxin transporter | | | 3.10 | 1.12 | | | | | | |
| | PA4144 | | probable outer membrane protein precursor | | | 3.33 | 1.15 | | | 2.44 | 1.22 | | |
| • | PA4205 | *mexG* | hypothetical protein | | | 43.50 | 1.21 | | | 6.82 | 1.18 | | |
| • | PA4206 | *mexH* | probable Resistance-Nodulation-Cell Division (RND) efflux membrane fusion protein precursor | | | 50.69 | 1.25 | | | 12.01 | 1.17 | | |
| • | PA4207 | *mexI* | probable Resistance-Nodulation-Cell Division (RND) efflux transporter | | | 36.89 | 1.23 | | | 10.22 | 1.18 | | |
| • | PA4208 | *opmD* | probable outer membrane protein precursor | | | 65.85 | 1.27 | | | 19.36 | 1.19 | | |
| • | PA4209 | *phzM* | probable phenazine-specific methyltransferase | | | 42.54 | 1.14 | | | 4.11 | 1.34 | 2.93 | 1.15 |
| • ○ | **PA4210** | ***phzA1*** | **probable phenazine biosynthesis protein** | **8.34** | **1.47** | **1788.72** | **1.26** | **6.27** | **1.49** | **13.02** | **2.11** | **63.05** | **1.33** |
| • ○ | **PA4211** | ***phzB1*** | **probable phenazine biosynthesis protein** | **23.71** | **1.34** | **1256.93** | **1.20** | **4.25** | **1.25** | **39.82** | **2.16** | **50.65** | **1.21** |
| ○ | PA4212 | *phzC1* | phenazine biosynthesis protein PhzC | | | 54.25 | 1.18 | | | 20.23 | 2.15 | 14.16 | 1.21 |
| ○ | PA4213 | *phzD1* | phenazine biosynthesis protein PhzD | | | 32.84 | 3.02 | | | 16.06 | 2.03 | 8.44 | 1.53 |
| • ○ | PA4214 | *phzE1* | phenazine biosynthesis protein PhzE | | | | | | | | | 6.89 | 1.23 |
| ○ | PA4215 | *phzF1* | probable phenazine biosynthesis protein | | | 208.87 | 1.25 | | | | | | |
| ○ | PA4216 | *phzG1* | probable pyridoxamine 5'-phosphate oxidase | | | 25.83 | 1.35 | | | | | | |
| • | PA4217 | *phzS* | flavin-containing monooxygenase | | | 18.47 | 1.22 | | | | | | |

| | | | | | | | | | | | | | | |
|---|---|---|---|---|---|---|---|---|---|---|---|---|---|---|
| ● | ○ | PA4306 | *flp* | Type IVb pilin, Flp | | | 2.72 | 1.23 | | | | | | |
| | | PA4348 | | conserved hypothetical protein | | | 2.36 | 1.19 | | | | | | |
| | | PA4384 | | hypothetical protein | | | 2.60 | 1.16 | | | | | | |
| | | PA4571 | | probable cytochrome c | | | 2.25 | 1.27 | | | | | | |
| | | PA4587 | *ccpR* | cytochrome c551 peroxidase precursor | | | 2.04 | 1.14 | | | | | | |
| | | PA4649 | *cupE2* | Pilin subunit CupE2 | | | 2.02 | 1.23 | | | | | | |
| | | PA4916 | *nrtR* | Nudix-related transcriptional regulator NrtR | | | 2.36 | 1.21 | | | | | | |
| | | PA5023 | | conserved hypothetical protein | | | 2.07 | 1.11 | | | | | | |
| | ○ | PA5162 | *rmlD* | dTDP-4-dehydrorhamnose reductase | | | | | | | 2.16 | 1.11 | | |
| | | PA5180 | | conserved hypothetical protein | | | 4.96 | 1.16 | | | | | | |
| | | PA5181 | | probable oxidoreductase | | | 5.16 | 1.16 | | | | | | |
| | | PA5219 | | hypothetical protein | | | 2.40 | 1.15 | | | 2.17 | 1.14 | | |
| ● | | PA5220 | | hypothetical protein | 2.70 | 1.10 | 15.23 | 1.16 | | | 6.49 | 1.23 | 3.33 | 1.11 |
| | | 2870682957 | | hypothetical protein | | | | | 2.57 | 1.08 | | | | |
| | | 2870698559 | | hypothetical protein | | | | | | | | | 4.15 | 1.10 |
| | | 2913673173 | | hypothetical protein | | | 4.95 | 1.25 | | | | | 2.02 | 1.22 |
| | | 2913675553 | | hypothetical protein | 3.14 | 1.18 | 4.78 | 1.17 | | | 2.79 | 1.26 | | |
| | | 2913676290 | | ectoine hydroxylase-related dioxygenase (phytanoyl-CoA dioxygenase family) | | | 2.54 | 1.16 | | | | | | |
| | | 2913679565 | | tRNA-dependent cyclodipeptide synthase | | | 85.20 | 1.30 | | | | | | |
| | | 2913679566 | | hypothetical protein | | | 8.44 | 1.34 | | | | | | |
| | | 2913679567 | | isopenicillin N synthase-like dioxygenase | | | 4.86 | 1.28 | | | | | | |
| | | 2913679568 | | cytochrome P450 | | | 3.32 | 1.24 | | | | | | |
| | | 2913679569 | | hypothetical protein | | | 5.10 | 1.29 | | | | | | |
| | | 2913679570 | | FIG00924305: hypothetical protein | | | 3.82 | 1.29 | | | | | | |
| | | 2913682544 | | hypothetical protein | 5.43 | 1.13 | | | | | 13.25 | 1.24 | 2.38 | 1.18 |
| | | 2913682924 | | hypothetical protein | 2.02 | 1.14 | | | | | | | | |
| | | 2913689358 | | hypothetical protein | | | | | | | 3.41 | 1.37 | | |
| | | 2913690790 | | transcriptional regulator with XRE-family HTH domain | | | | | | | 5.70 | 1.14 | | |
| | | 2913690791 | | integrating conjugative element relaxase (TIGR03760 family) | | | | | | | 2.28 | 1.15 | | |
| | | 287.13131.peg.236 | | hypothetical protein | | | 2.64 | 1.16 | | | | | | |
| | | 287.13131.peg.2743 | | hypothetical protein | | | 6.51 | 1.62 | | | | | | |
| | | 287.13131.peg.2917 | | hypothetical protein | | | 7.06 | 1.83 | | | | | | |
| | | 287.13131.peg.4268 | | hypothetical protein | | | 33.68 | 1.18 | | | | | | |
| | | 287.13131.peg.6440 | | FIG00957537: hypothetical protein | | | 2.49 | 1.31 | | | 8.64 | 1.45 | 3.58 | 1.30 |
| | | 287.13131.peg.793 | | FIG00958200: hypothetical protein | | | 184.48 | 1.21 | 3.10 | 1.37 | | | | |
| | | 287.13133.peg.4085 | | FIG00955856: hypothetical protein | | | | | | | 7.14 | 1.27 | | |
| | | 287.13133.peg.4120 | | hypothetical protein | | | | | | | 5.98 | 1.39 | | |
| | | 287.13133.peg.5885 | | hypothetical protein | | | 5.29 | 1.25 | | | 4.34 | 1.30 | | |
| | | 287.9771.peg.292 | | hypothetical protein | | | | | 2.44 | 1.10 | | | | |

[a]Locus tags, names, and functions from the PAO1 annotation on Pseudomonas.com, with the following exceptions: for genes not present in PAO1, the IMG gene ID and function are used; for genes not present in PAO1 or IMG annotations, RAST gene ID and function are used; *aze* annotation from Patteson *et al.* 2019. ●, gene was reported as QS-regulated in PAO1 in Chugani *et al.* 2012. ○, gene shows evidence for direct transcriptional regulation by RhlR, or first gene in operon shows such evidence. **Bold** indicates genes in the core RhlR regulon reported in this study (Table 2, main text).

[b]Fold-change from individual comparisons between wild-type CF isolates and their isogenic *rhlR* mutants. FC, fold-change; FCSE, fold-change standard error.

**Table S5. Bacterial strains and plasmids.**

| Strain or plasmid | Relevant properties | Reference or origin |
|---|---|---|
| *Pseudomonas aeruginosa* | | |
| PAO1 | Wild-type laboratory strain | (Stover *et al.,* 2000) |
| PAO *lasR* | PAO1 derivative; "DA5" unmarked in-frame *lasR* deletion mutant | (Siehnel *et al.*, 2010) |
| PAO *lasR rhlR* | PAO1 derivative; "DA6" unmarked double-null deletion mutant in which both *lasR* and *rhlR* harbor in-frame deletions: '*lasR rhlR*' | (Siehnel *et al.*, 2010) |
| E104 | Wild-type cystic fibrosis isolate | (Feltner *et al.*, 2016) |
| E104 *rhlR* | E104 derivative; unmarked in-frame *rhlR* deletion from residue 3-240 | This study |
| E113 | Wild-type cystic fibrosis isolate | (Feltner *et al.*, 2016) |
| E113 *rhlR* | E113 derivative; unmarked in-frame *rhlR* deletion from residue 3-240 | This study |
| E125 | Wild-type cystic fibrosis isolate | (Feltner *et al.*, 2016) |
| E125 *rhlR* | E125 derivative; unmarked in-frame *rhlR* deletion from residue 3-240 | This study |
| E131 | Wild-type cystic fibrosis isolate | (Feltner *et al.*, 2016) |
| E131 *rhlR* | E131 derivative; unmarked in-frame *rhlR* deletion from residue 3-240 | This study |
| E167 | Wild-type cystic fibrosis isolate | (Feltner *et al.*, 2016) |
| E167 *rhlR* | E167 derivative; unmarked in-frame *rhlR* deletion from residue 3-240 | This study |
| *Escherichia coli* | | |
| DH5α | F- Φ80lacZYA-argF U169 recA1 hsdR17 (rk-, mk+) phoA supE44 λ- thi-1 gyrA96 relA1 | Invitrogen |
| S17-1 | *recA* pro *hsdR* RP4-2Tc::Mu-Km::Tn7 | (Simon *et al.,* 1983) |
| *Plasmids* | | |
| pEXG2 | Conjugative suicide plasmid for allelic exchange; $Gm^R$, *sacB* | (Hmelo *et al.,* 2015) |
| pEXG2-E104rhlR-KO | pEXG2 containing an E104 strain-specific in-frame deletion allele of *rhlR* from amino acid 3-240 | This study |
| pEXG2-E113rhlR-KO | pEXG2 containing an E113 strain-specific in-frame deletion allele of *rhlR* from amino acid 3-240 | This study |
| pEXG2-E125rhlR-KO | pEXG2 containing an E125 strain-specific in-frame deletion allele of *rhlR* from amino acid 3-240 | This study |
| pEXG2-E131rhlR-KO | pEXG2 containing an E131 strain-specific in-frame deletion allele of *rhlR* from amino acid 3-240 | This study |
| pEXG2-E167rhlR-KO | pEXG2 containing an E167 strain-specific in-frame deletion allele of *rhlR* from amino acid 3-240 | This study |
| pProbe-GT | Broad-host-range vector with a promoterless *gfp*, $Gm^R$ | (Miller *et al.*, 2000) |
| pProbe-GT-$P_{lasI}$-*gfp* | pProbe-GT with *gfp* under the control of the *lasI* promoter | (Feltner *et al.*, 2016) |
| pBBR-*gfp* | Promoterless *gfp* transcriptional reporter, $Gm^R$ | (Smalley *et al.*, 2021) |
| pBBR-$P_{rhlA}$-*gfp* | pBBR-*gfp* with *gfp* under the control of the *rhlA* promoter | This study |

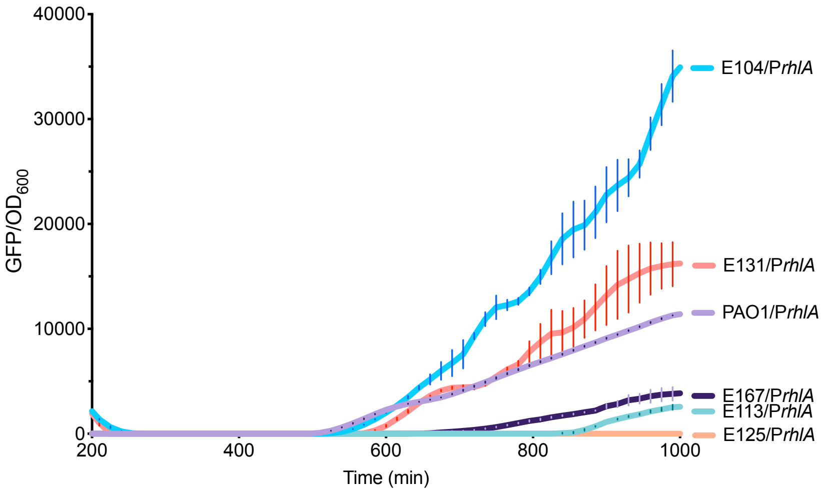


**Figure S1. RhlR-specific transcriptional activity.** pP$_{rhlA}$-*gfp* reporter activity, normalized to $OD_{600}$ and corrected for background fluorescence using a promoterless *gfp* control plasmid. PAO1 *lasR* was also tested, but yielded values of zero at all timepoints, and so is omitted from the graph.

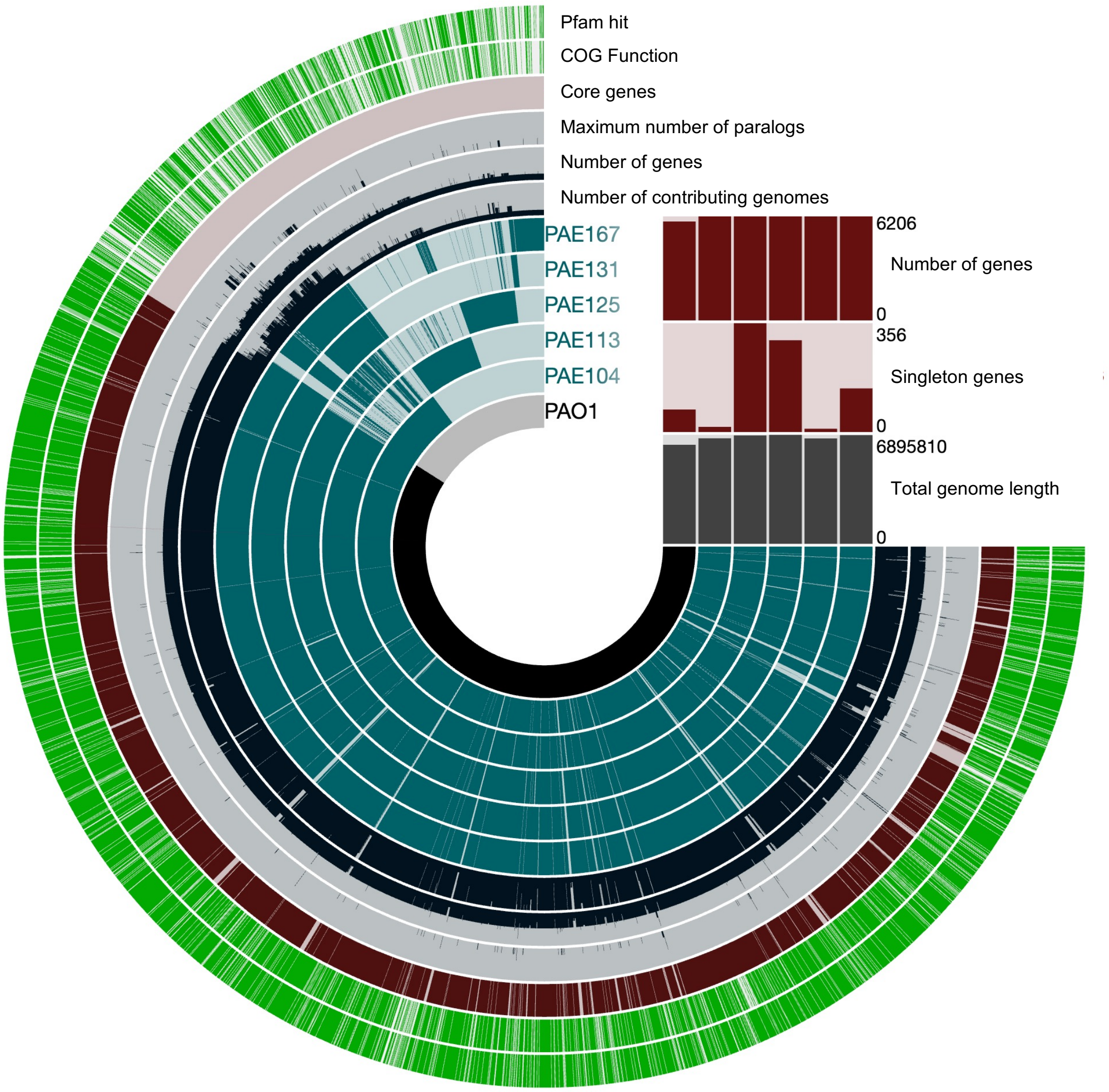


**Figure S2. Summary figure of pangenomic analysis.** Genes are indicated as the presence of a band in each circular track, and ordered by forced synteny to PAO1. Bands for Number of contributing genomes, Number of genes, and Maximum number of paralogs are continuous variables. The outer two tracks indicate a significant Pfam or COG database hit during computation in the anvi'o software ecosystem[1].

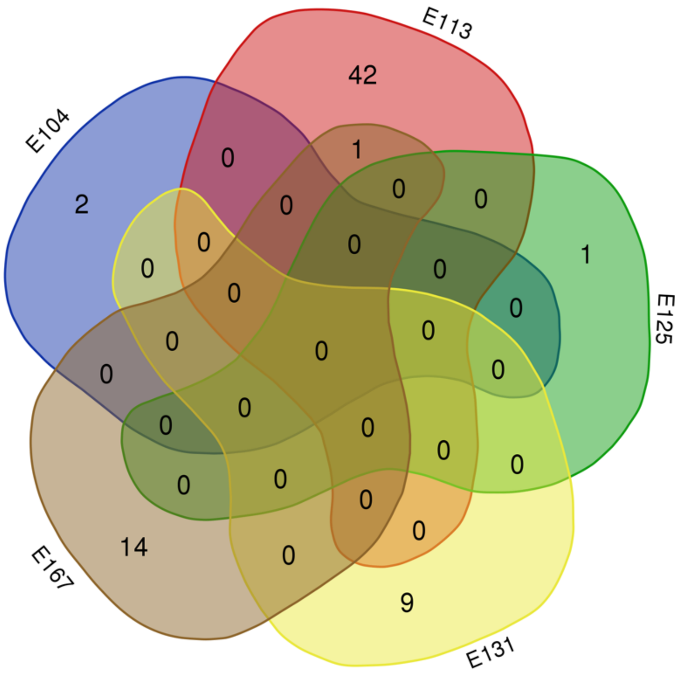


**Figure S3. RhlR repressed genes: core analysis.** Venn diagram displaying overlap of RhlR repressed genes in individual CF isolate regulons. Lobes not scaled to size.
